# Loss of TP53 mediates suppression of Macrophage Effector Function via Extracellular Vesicles and PDL1 towards Resistance against Chemoimmunotherapy in B-cell malignancies

**DOI:** 10.1101/2020.06.11.145268

**Authors:** Elena Izquierdo, Daniela Vorholt, Benedict Sackey, Janica L. Nolte, Stuart Blakemore, Jan Schmitz, Verena Barbarino, Nadine Nickel, Daniel Bachurski, Ludmila Lobastova, Milos Nikolic, Michael Michalik, Reinhild Brinker, Olaf Merkel, René Neuhaus, Maximilian Koch, Gero Knittel, Lukas Frenzel, Hans Christian Reinhardt, Martin Peifer, Rocio Rebollido-Rios, Heiko Bruns, Marcus Krüger, Michael Hallek, Christian Pallasch

## Abstract

Chemoimmunotherapy (CIT) is the standard of care in B-cell malignancies. It is relying on synergistic effects of alkylating chemotherapy and monoclonal antibodies via secretory crosstalk with effector macrophages. Here, we observed that loss of p53 function mediates resistance to CIT by suppressing macrophage phagocytic function.

Loss of p53 leads to an upregulation of PDL1 and an increased formation of extracellular vesicles (EVs). EVs directly inhibit macrophage phagocytosis by PDL1 surface expression. Suppression of phagocytic function by lymphoma cell-derived EVs could be abrogated by pre-incubation of EVs with anti-PDL1 antibodies, CRISPR-KO of *PDL1* and abrogation of EV formation by *RAB27A*-KO in lymphoma cells. Immune checkpoint inhibition represents a viable strategy to overcome EV-mediated resistance to chemoimmunotherapy in lymphoma.

**Significance:** Loss of *TP53* mediates cell autonomous resistance to genotoxic chemotherapy, moreover non-cell autonomous effects may cause therapy resistance mediated by the tumor microenvironment. We identify a *TP53*-dependent mechanism that mediates resistance to synergistic chemoimmunotherapy by increasing formation of EVs and expression of the PDL1 immune checkpoint. PDL1 on EVs is directly responsible for macrophage suppression, preventing the exertion of the essential effector function of antibody-dependent cellular phagocytosis. This novel mechanism of resistance is in turn targetable by PDL1 checkpoint inhibition. Enhanced EV-release and immune checkpoint expression in lymphoma are novel mechanisms of macrophage modulation in the lymphoma microenvironment. We provide a novel principle of resistance to chemoimmunotherapy (CIT) representing of immediate relevance to treatment of refractory B-cell lymphoma.

**Highlights:**

- Loss of *TP53* in B-cell lymphoma induces resistance towards chemoimmunotherapy (CIT) by inhibition of macrophage effector function through PDL1 upregulation
- Loss of *TP53* increases formation of extracellular vesicles (EVs) carrying PDL1
- EVs inhibit antibody-mediated cellular phagocytosis (ADCP), a key macrophage effector function in CIT
- Targeting PDL1 on EVs with immune checkpoint inhibitors overcomes *TP53*-mediated resistance to CIT

## Introduction

The tumor microenvironment is characterized by multiple reciprocal interactions of malignant cells with non-malignant stroma or immune cells (Spranger and Gajewski, 2018). Particularly, macrophages are at center stage in this network, determining disease progression, therapeutic response, as well as refractory niches (Hughes *et al*., 2015; Lux *et al*., 2014; Pallasch *et al*., 2014a; Qian and Pollard, 2010). In previous work, we and others could show that macrophages exert antibody-dependent cellular phagocytosis (ADCP) and represent the essential mediator of synergy in the administration of the chemoimmunotherapy (CIT) of aggressive B cell lymphoma(Lossos et al., 2019; Pallasch et al., 2014a; Roghanian et al., 2019). This specific combination treatment strongly increases tumor clearance by repolarization of tumor-associated macrophages from a suppressed state to an activated phenotype (Lossos et al., 2019; Pallasch et al., 2014b). CIT serves as a standard therapy in numerous B-cell malignancies and is most frequently applied by anti-CD20 antibodies, such as rituximab in combination with either Fludarabine/Cyclophosphamide (R-FC) or Cyclophosphamide, Doxorubicin, Vincristine and Prednisone (R-CHOP) (Coiffier *et al*., 2002; Hallek *et al*., 2010). Despite the success of CIT in the front-line setting, therapy of relapsed or refractory disease particularly in diffuse large B cell lymphoma (DLBCL) still imposes a major clinical challenge. Although distinction between GCB- and ABC subtypes and further molecular subclassification opened avenues for tailored treatment strategies, limited molecular markers are available to tailor frontline therapy to overcome primary resistant lymphoma (Chapuy *et al*., 2018; Wilson *et al*., 2015). Notably, mutations of *TP53* are present in approximately 20% of DLBCL; thus, representing a significant subgroup of patients (Cancer Genome Atlas Research Network *et al*., 2013; Chapuy *et al*., 2018; Lohr *et al*., 2012; Zhang *et al*., 2013).

Loss or mutation of *TP53* has been identified as an important mediator of chemoresistance in a variety of malignant entities, due to its central coordinating function in multiple cellular stress responses (Mantovani et al., 2019). Moreover, loss of *TP53* mediates pro-tumorigenic alterations in the tumor microenvironment (Cooks *et al*., 2013, 2018). It remains to be clarified how alterations of the DNA damage pathway in malignant cells functionally affect the complex interactions and outcome of CIT.

Here, we primarily approached disruptions in the DDR cascade affecting the synergistic effects of CIT. Our data indicate that particularly *TP53* functional status determines phagocytic function and macrophage-dependent therapeutic response to monoclonal antibodies. As a novel mediator of this interaction, we identify increased EV release and PDL1 immune checkpoint expression, due to loss of *TP53* function in lymphoma.

## Results

### Efficient phagocytosis in chemoimmunotherapy response of B-cell malignancies depends on DNA damage response pathway integrity

To dissect the mechanism of the synergistic interaction between alkylating chemotherapy and monoclonal antibodies (Leskov *et al*., 2013; Lossos *et al*., 2019; Pallasch *et al*., 2014a), we disrupted key components of the DDR pathway in two different aggressive B-cell lymphoma models: the hMB humanized Double-Hit-Lymphoma model (Leskov *et al*., 2013) and the *Myd88* p.L252P-driven DLBCL model *(Knittel et al*., 2016). In both cases, we generated shRNA-mediated knockdowns of genes that code for the key DDR mediators: CHK1, CHK2, DNAPK (*PRKDC*), MK2 (*MAPKAPK2*), p38-alpha, ATM, DNA-PK and p53. We exposed shRNA-defined DDR component-deficient cells to mafosfamide, an *in vitro* alkylating agent as surrogate for cyclophosphamide effects. We were then determining the antibody-dependent cellular phagocytosis (ADCP) using the anti-CD52 antibody alemtuzumab as a proof of principle therapeutic antibody in co-culture assays with J774A.1 macrophages (Figure 1A). In line with previous reports (Leskov et al., 2013; Lossos et al., 2019; Pallasch et al., 2014a; Roghanian et al., 2019), we observed that chemotherapy pretreatment of hMB lymphoma B-cells significantly enhanced their susceptibility for phagocytic engulfment, as compared to untreated hMB cells (Figure 1A-B). However, disruption of the DDR led to an abrogation of this phagocytosis enhancement in *CHK1, CHK2, PRKDC, MAPKAPK2, p38a, ATM, DNA-PK* and *TP53*-shRNA targeted knockdown hMB cells (Figure 1B). Since we and others have previously identified a secretory crosstalk between lymphoma B-cells and macrophages as a partially responsible mechanism for the increase of tumor clearance (Lossos *et al*., 2019; Pallasch *et al*., 2014a; Roghanian *et al*., 2019), we next aimed to address the influence of the DDR-induced acute secretory response on macrophage phagocytic capacity. We generated conditioned media from mafosfamide-treated or untreated hMB control cells and the DDR shRNA-targeted cells, respectively (Figure 1C). We confirmed that conditioned media obtained from shRNA-control empty vector (*shCTRL*) hMB cells exposed to mafosfamide significantly enhanced phagocytic capacity of macrophages, in comparison to conditioned media from untreated cells (Figure 1D). However, the stimulatory effect of conditioned media was not detected with the media generated from mafosfamide-treated *CHK1, CHK2, PRKDC, MAPKAPK2, p38a, ATM*, and *TP53*-shRNA targeted hMB cells (Figure 1D). To employ an independent model of aggressive lymphoma, we utilized the *Myd88* p.L252P-driven DLBCL model (Figure 1F-H). First, we observed that combination treatment of mafosfamide with the monoclonal antibody anti-CD20 (18B12) improved the phagocytosis of *shCTRL Myd88* p.L252P cells by primary peritoneal and bone marrow-derived macrophages, as well as by the macrophage cell line J774A.1 (Figure S1C). Subsequently, we compared the ADCP of shRNA defined lymphoma cells and observed that in the absence of chemotherapy treatment, lymphoma cells deficient for *Mapkapk2, Chk2* and *p38* showed resistance to phagocytosis in treatment-free co-culture conditions and also in the presence of anti-CD20 antibody (Figure 1E,F). When applying alkylating treatment to co-cultures, *Chk1*-, *Prkdc* and *Mapkapk2*-deficient lymphoma cells were sensitized to therapeutic effects. Interestingly, *Tp53*-deficient cells displayed the highest resistance towards chemoimmunotherapeutic combination (Figure 1F, G). Altogether, alterations in the DDR pathway in lymphoma B-cells cells affect macrophage anti-tumor features and particularly alterations of *TP53* provide resistance to combinations of alkylating chemotherapy and therapeutic antibodies.

**Figure 1.**
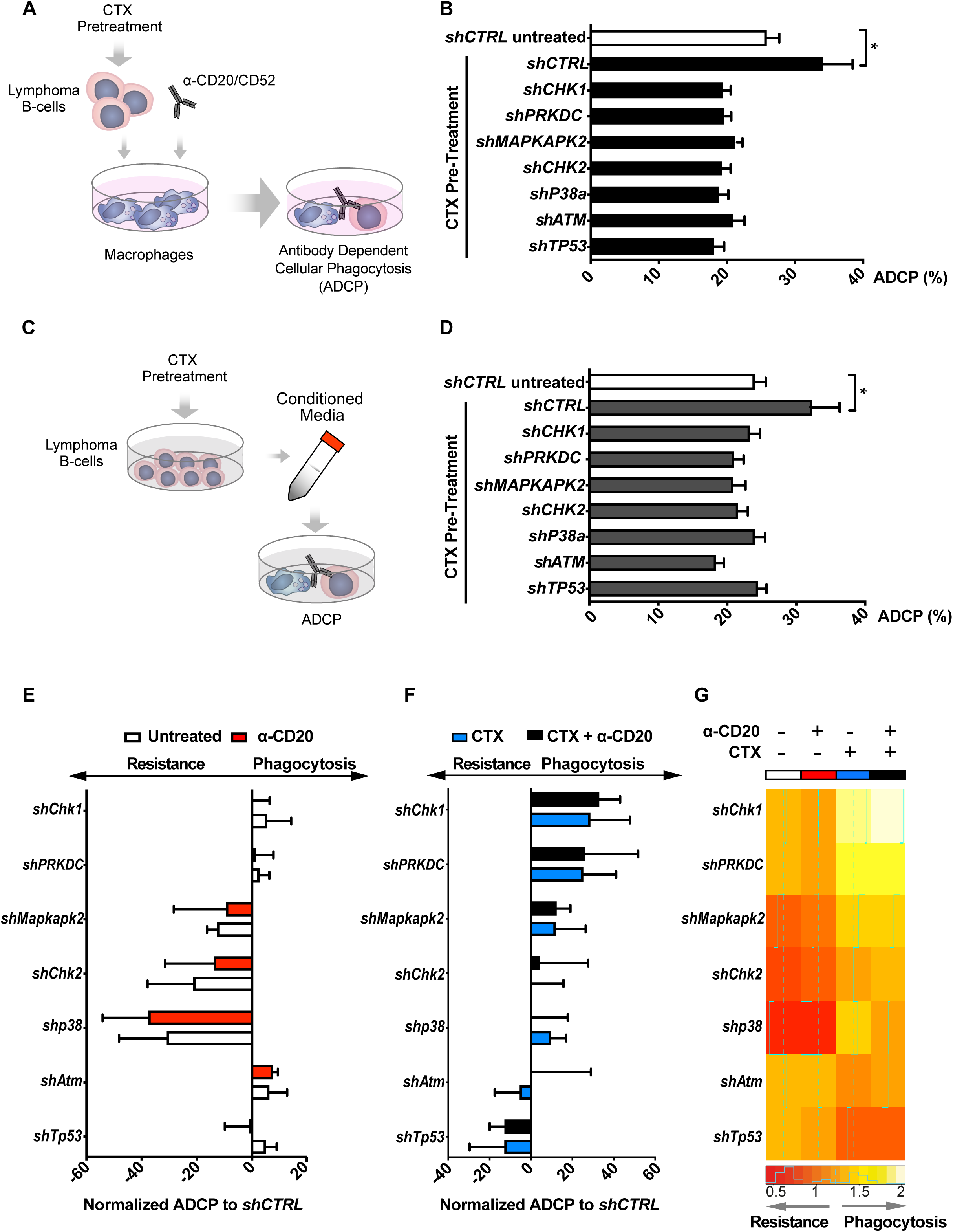
CIT-induced antibody dependent cellular phagocytosis (ADCP) of lymphoma B-cells with altered DNA damage response signaling pathway. **A)** Schematic representation of an ADCP assay. **B)** Alemtuzumab (anti-CD52)-mediated ADCP of empty vector (*shCTRL*) and the indicated DDR-knockdown hMB cells, pre-treated or not with CTX (3µM) and co-cultured with peritoneal macrophages. **C)** Schematic representation of an ADCP assay in the presence of conditioned media.**D)** Alemtuzumab-mediated ADCP of *shCTRL* hMB cells in the presence of conditioned media generated from CTX-treated hMB-DDR knockdown cells or *shCTRL* cells and co-cultured with peritoneal macrophages. **F-H)** ADCP of the indicated DDR-knockdown Myd88p.L252P cells normalized to the *shCTRL* cells after: **F)** treatment or not with anti-CD20 antibody (18B12) at 50µg/ml, and **G)** treatment with CTX (1.64 µM) alone or in combination with anti-CD20 antibody. **H)** Heatmap representation of the phagocytosis pattern of Myd88p.L252P cells co-cultured with peritoneal macrophages under different treatment conditions. Broken line represents *shCTRL*cells and the solid stepped lines indicate quantitatively, average phagocytosis of the shRNA-targeted cells normalized to *shCTRL* for the indicated gene (n=5). All the graphics showed the mean ± SD of at least 3 independent experiments. (\**p*□<□0.05).

### Macrophage phagocytic capacity upon chemotherapy is diminished in *TP53-*aberrated patient samples *in vitro*, and in the Eµ-*TCL1/Tp53*^*fl/fl*^ mouse model *in vivo*

In view of the influence of the DDR pathway on the generation of an adequate response to CIT, we aimed to elucidate the role of p53 as a regulatory node, coordinating the interaction of transformed B-cells with the microenvironment. Triggering DDR by alkylating treatment, p53 protein expression was diminished in *CHK1*-, *CHK2*-, *TP53*- and *PRKDC*-shRNA targeted hMB cells (Figure S1A). Thus, we speculated that the disruption of CIT synergy response when altering DDR components could be a result of insufficient p53 activation. In order to evaluate the effects of p53 activation in the absence of DNA damage, we utilized nutlin-3A as an indirect activator of p53. Interestingly, nutlin-3A similarly modified the ADCP of control and *CHK1*-, *CHK*2-, *p38a*- and *MAPKAPK2*-deficient hMB cells, but left *TP53*-deficient lymphoma cells unaffected, which exhibited a significantly reduced phagocytosis level (Figure 2A). In the same way, nutlin-3A treatment of *PRKDC*- and *ATM*-deficient cells did not alter macrophage phagocytic capacity, which correlates with the absence of p53 induction by nutlin-3A (Figure S2B). Altogether, these results indicate that p53 function in lymphoma B-cells influences the response of macrophages to CIT.

**Figure 2.**
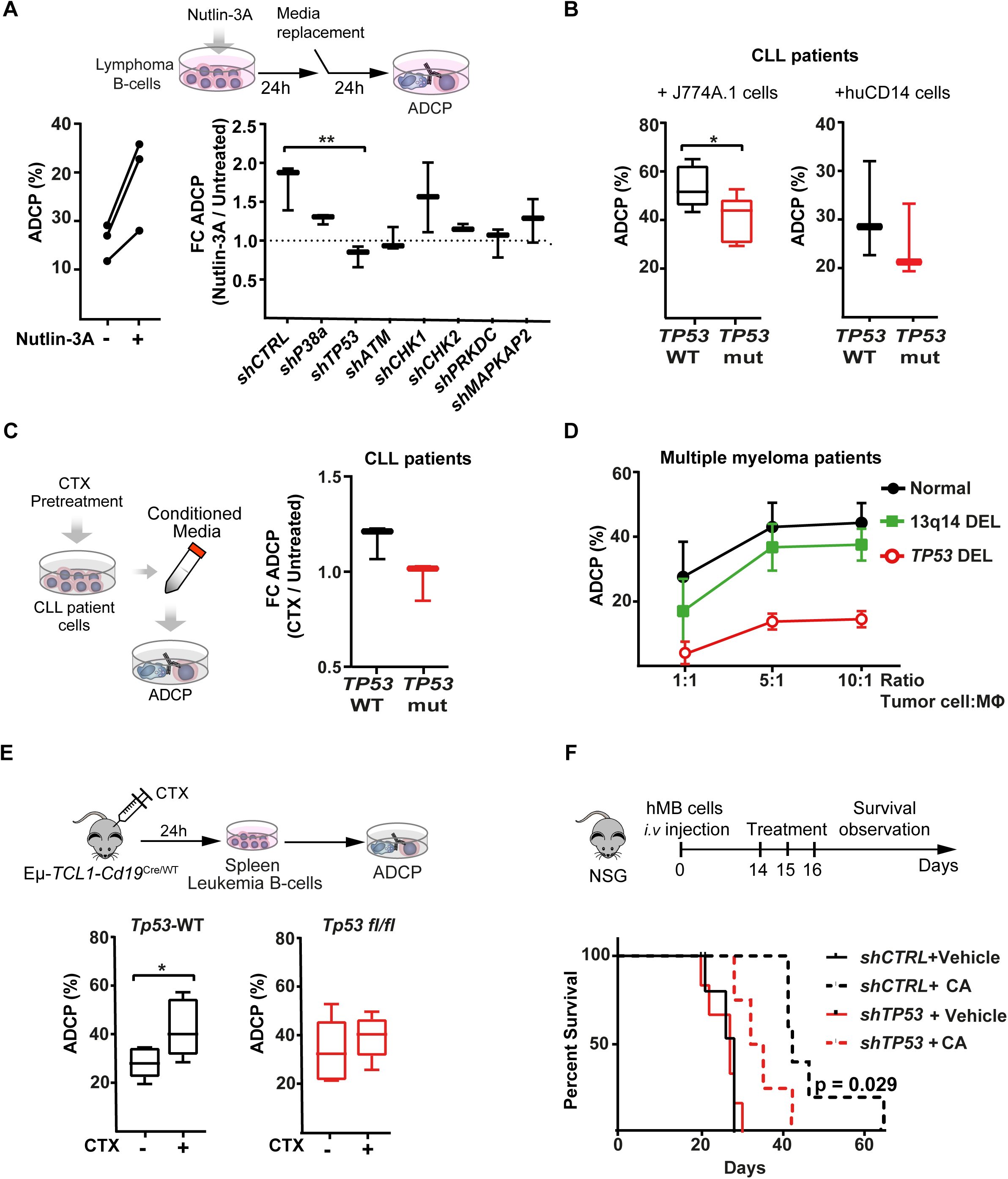
Effect of *TP53* loss in the CIT response of lymphoma malignances. **A-C)** Alemtuzumab-mediated ADCP of:*shCTRL* **A)** hMB cells, pretreated or not with nutlin-3A (left panel) (n=3). Right panel shows ADCP ratio of pretreated nutlin-3A vs untreated empty vector and the indicated DDR-knockdown hMB cells (n=3); **B)** CLL cells from patients with *TP53* wildtype (WT) and mutant expression (*TP53* mut) on the presence of murine J774A.1 (n=6) or human CD14^+^-derived macrophages (n=2); **C)** CLL cells in the presence of conditioned media generated from CLL cells treated or not with mafosfamide (CTX). Left panel shows a schematic representation of the approach and right panel the fold change on the ADCP of WT and TP53 mutant CLL cells, after incubating the cells with mafosfamide-conditioned media (n=3). **D)** Daratumumab (anti-CD38 antibody)-mediated ADCP of multiple myeloma tumor cells obtained from patients with normal *TP53* expression (normal karyotype or 13q14 deletion) and from patients with *TP53* deletion. In all cases the tumor cells were co-cultured with autologous macrophages in the indicated ratios (n=3). **E)** TCL1/wt *and* TCL1/*Tp53*^*fl/fl*^ leukemic mice were treated or not with 10 mg/kg cyclophosphamide (CTX) for 24 h. Then, the spleen leukemic cells were isolated and used to perform an alemtuzumab-mediated ADCP in the presence of peritoneal macrophages (n=3). **F)** NSG mice were *i*.*v* injected with shRNA-control (*shCTRL*) or shRNA-*TP53* (*shTP53*) hMB tumor cells. After 10 days, mice were treated *i*.*p* with cyclophosphamide and alemtuzumab (CA), as CIT combination, or PBS as control treatment. (n=5-6). (\**p*□<□0.05).

To further confirm our observation, we used primary chronic lymphocytic leukemia (CLL) patient samples defined by their *TP53* status (Figure 2B). Notably, macrophages co-cultured with mafosfamide pre-treated *TP53*-mutant CLL cells in the presence of alemtuzumab showed significantly lower ADCP than macrophages co-cultured with *TP53-*wild type cells (Figure 2B). This finding was corroborated, using an independent source of macrophages, including human primary CD14^+^ differentiated monocytes (Figure 2B). In addition, the effect of the CLL-conditioned media on macrophage phagocytic capacity upon chemotherapy was evaluated (Figure 2C). Here, we could observe an increase in tumor clearance after the addition of mafosfamide-treated media derived from *TP53*-wildtype CLL cells, whereas this stimulatory effect was absent in *TP53*-mutant CLL-derived conditioned media. Likewise, the effect of *TP53* on tumor cell-macrophage crosstalk was investigated using primary malignant multiple myeloma patient cells, co-cultured with autologous macrophages in the presence of the anti-CD38 antibody daratumumab (Figure 2D)(Overdijk *et al*., 2012). Our data showed that macrophages were able to phagocytose approx. 50% of myeloma cells from patients with normal karyotype del13q14-aberration and wildtype *TP53*. Conversely, *TP53*-deficient patients displayed a significantly reduced treatment response of 6.4% phagocytosis even in high 10:1 effector-to target cell ratio (Figure 2D)(Busch et al., 2018). Conversely, *TP53*-deficient patients displayed a significantly reduced treatment response of 6.4% phagocytosis, even in high 10:1 effector-to target cell ratio (Figure 2D). To validate the relevance of *TP53* as a central regulator of microenvironment-dependent treatment response *in vivo*, we next utilized the *Eµ-TCL1* mouse model of CLL. Here, leukemic *Eµ-TCL1*/*Cd19*-*Tp53* wildtype (TCL1/wt) and *Eµ-TCL1/Cd19*^*Cre/wt*^*-Tp53* ^*fl/fl*^ (TCL1/*Tp53*^*fl/fl*^) mice (Knittel *et al*., 2017) were treated with cyclophosphamide. After 24 hours leukemic cells were obtained for *ex vivo* ADCP assessment targeted by the murine-specific anti-CD20 antibody 18B12. In line with our previous observations, *in vivo* chemotherapy treatment significantly improved the ADCP of TCL1/wt*-*derived leukemia cells, while we could not detect an increase of therapy response in TCL1/*Tp53*^*fl/fl*^-derived cells (Figure 2E). These results indicate that loss of *TP53* switches malignant B cells towards a refractory state against CIT. To verify our hypothesis, we studied the *in vivo* response to CIT with the humanized mouse model of Double-hit lymphoma. Specifically, *shTP53*-transduced hMB alongside *shCTRL* cells were transplanted into immunodeficient NSG recipient mice and CIT treatment (cyclophosphamide and alemtuzumab; CA) was initiated, upon disease onset (Figure 2F). CIT resulted in a significantly increased survival of mice transplanted with *shCTRL*-hMB cells (CA median = 40 ± 1.73 days vs. PBS median = 27 ± 1.22 days, p = 0.029). However, the response of mice bearing the *shTP53*-hMB cells to the combination therapy showed a significantly shorter overall survival (CA median = 28 ± 1.65 days vs. PBS median = 28 ± 0.601 days, p = 0.125), compared to *shCTRL*-hMB cells (Figure 2F). In summary, *TP53* expression serves as a central regulatory node for interactions with macrophages in the TME, particularly in determining response to CIT in B-cell malignancies..

### *Tp53* deficiency in leukemic B-cells represses the expression of a chemotherapy-induced phagocytosis gene signature in TME macrophages

Since macrophages are extremely sensitive to the surrounding extracellular milieu, we asked whether macrophage dysfunction in states of *Tp53*-deficient tumors was due to a switch in the secretory response of lymphoma B-cells. However, we could not observe p53-dependent changes causative to the above-described resistant phenotype (Figure S3A, B). To further understand the p53-dependent alterations towards communication between macrophages and malignant B-cells, we investigated the *in vivo* effect of chemotherapy on the TME by single-cell transcriptomic analysis. Briefly, TCL1/wt-mice and TCL1/*Tp53*^*fl/fl*^ were injected with cyclophosphamide. CD19-negative TME cells were obtained after 24 hours from the spleen and subjected to single-cell RNA sequencing (scRNA-seq) analysis (Figure 3A, upper panel). After raw data quality control and read count data pre-processing, we ended up with a dataset of 21,782 cells across all conditions (Figure 3A, lower panel). Using Uniform Manifold Approximation and Projection (UMAP) dimension reduction (Figure 3A, middle panel), we could observe 18 cell clusters, representing 8 main cell populations, from which macrophages were the second highest represented after T-cells (Figure S3D, Table S1.1). Overall, we found a larger proportion of macrophages in the TCL1/*Tp53*^*fl/fl*^ mice, than in the control TCL1/wt-mice (Figure 3B). Chemotherapy treatment generally reduced the number of macrophages in both, TCL1/*Tp53*^*fl/fl*^ and TCL1/wt TME (Figure 3B) and mainly decreased the number of significant genes affected on TCL1/*Tp53*^*fl/fl*^-derived macrophages (Suppl. Table S1.2). Specifically, macrophage cluster 2 (MΦ-C2) showed pronounced gene ratio differences upon chemotherapy, which gene enrichment analysis (p-adjust ≤ 0.05) displayed GO terms associated to phagocytosis and cytokine function (Figure 3D). Likewise, our data showed that chemotherapy treatment downregulated the phagocytic, cytokine and myeloid-related gene expression profile of TCL1/*Tp53*^*fl/fl*^-mice macrophages, while having the opposite effect on macrophages obtained from TCL1/wt TME (Figure 3E and Figure S3E, Table S1.3). Thus, in the same line as our *in vitro* observation, we confirmed that loss of *TP53* in malignant B-cells similarly affects their communication with macrophages in the TME *in vivo*, mainly altering macrophage phagocytic capacity and immune response.

**Figure 3.**
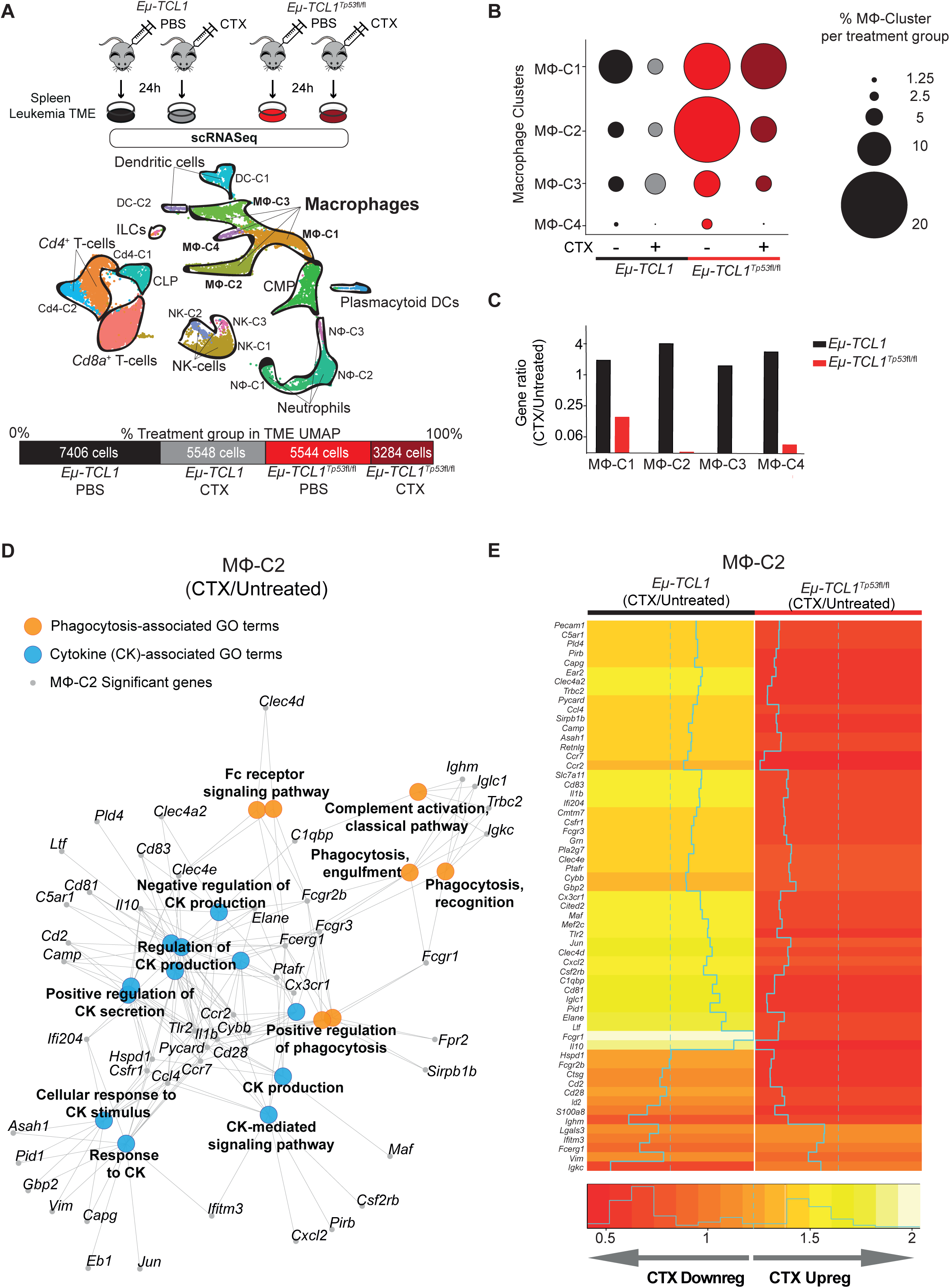
Characterization of TME-macrophage properties upon chemotherapy on *Eμ-TCL1* mice. **A-E** TCL1/wt *and* TCL1/*Tp53*^*fl/fl*^ leukemic mice were *i*.*p* injected 10 mg/kg cyclophosphamide (CTX) or PBS for 24 h (1 mouse per treatment). **A)** Schematic representation of the experimental design (upper panel). TME CD19-negative cells were isolated from the spleens of each treatment group, followed by single-cell RNA sequencing (scRNA-seq). Integrated Uniform Manifold Approximation and Projection (UMAP) dimension reduction plot of all treatment groups (n = 21,782 cells) (centre panel). Cells are colored by the clusters determined by cell type. Percentage and total cell number of each treatment group contributing to the integrated UMAP (lower panel). **B)** Dot plot representing the percentage of the different identified macrophage clusters (MF-C1, MF-C2, MF-C3 and MF-C4) within the TME of each treatment group in relation to the total number of macrophages of each cluster. **C)** Bar plot showing the ratio of significant genes observed in every macrophage cluster upon chemotherapy versus untreated condition. **D)** Representation of selected GO terms enriched in the MF-C2 and the genes of the GO term. **E)** Heatmap showing fold change gene expression induced by CTX of significant genes of the MΦ-C2.

### Chemotherapy treatment modifies the proteomic profile of lymphoma B-cells in a p53dependent manner and indicates altered extracellular vesicle formation

We have clearly identified loss of *TP53* a mechanism to enable lymphoma cells to inactivate macrophages. The use of conditioned media from respective cells indicates soluble mediators of this effect. However, using a multiplex cytokine assessment we could not reveal any TP53 specific effects as an explanation. Therefor the proteomic profile of *shCTRL* and *shTP53* lymphoma B-cells after mafosfamide treatment was next analysed by a mass spectrometry label-free protein quantification approach (Figure 4A). In total, more than 4000 proteins were detected in every condition (Table S2.1). Hierarchical agglomerative clustering clearly differentiated *shCTRL* from *shTP53* proteome (Figure S4A). To a minor extent, mafosfamide exposure for 6 h significantly (p-adjust ≤ 0.05; 1≤ log_2_ fold change ≤-1) affected the protein expression profile of control and *TP53*-depleted lymphoma B-cells (Figure S4A-D). Our results showed that *TP53* depletion profoundly influenced the proteomic profile of lymphoma B-cells, showing a downregulation of 36 proteins and the upregulation of another 60, compared to the control cells (Figure 4A). Interestingly, we observed that mafosfamide treatment differently modified the proteomic profile of *TP53*-deficient and control cells (Figure S4C, D). Mafosfamide treatment significantly regulated the expression of 101 proteins on *TP53*-deficient cells, compared to control cells and produced a shift in the number of up- and downregulated proteins controlled by *TP53* (Figure 4A). Analyzing GO terms of significantly (p-adjust ≤ 0.1) regulated proteins (Table S2.2), we observed that loss of *TP53* expression mainly correlated to changes on actin cytoskeleton, extracellular plasma membrane and vesicle-related terms, composed of proteins such as CD9, vacuolar protein sorting 33A (VPS33A), and sorting nexin 9 (SNX9) (Figure 4B, C). Similarly, mafosfamide treatment of *shTP53* cells differently regulated proteins associated to actin cytoskeleton and cell adherens junction processes, displaying altered expression levels of extracellular vesicle (EV)-associated proteins, such as annexins 1 and 6 (ANXA1 and ANXA6) and the tetraspanin CD81 (Figure 4B, D). Therefore, proteomic analysis has indicated that *TP53* loss in lymphoma B-cells involves actin cytoskeleton reorganization and alterations in extracellular vesicle formation, which we hypothesize, could be responsible for the communication between the malignant B-cells and the TME macrophages.

**Figure 4.**
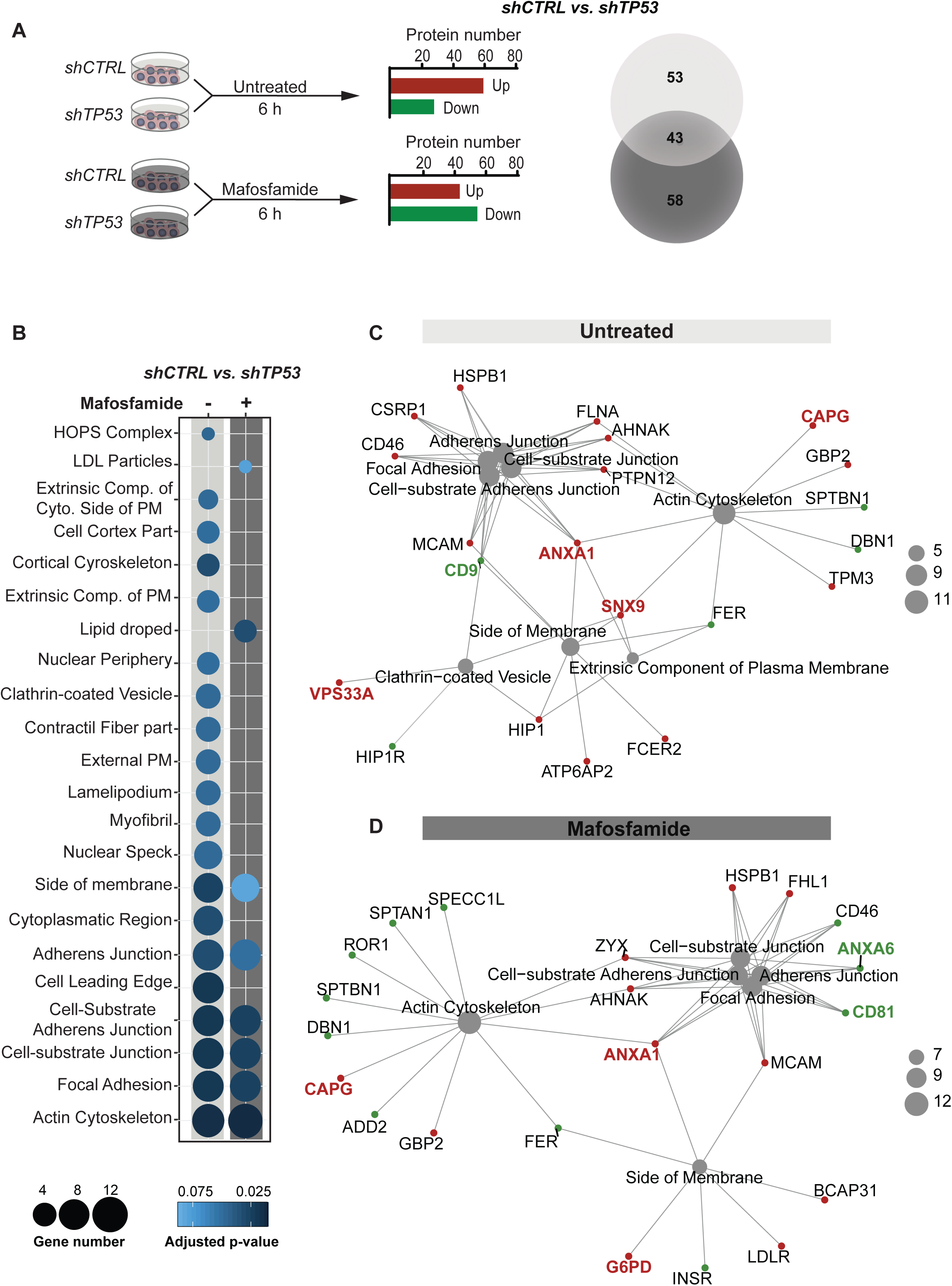
Chemotherapy-rewiring of control and *TP53*-deficient lymphoma B-cells proteome. **A-D)** *shCTRL* and *shTP53* hMB cells were treated with 3µM mafosfamide or vehicle (untreated) for 6 h and their proteomic profile were analyzed by mass spectrometry label-free protein quantification (n=3). **A)** Workflow of the experiment (left panel). Bar plot of the number of up- and downregulated proteins that are differentially expressed in *shTP53* hMB cells in comparison to *shCTRL* hMB cells, either in untreated or mafosfamide-treated condition (central panel). Venn diagram illustrating the total number of significant proteins specifically or commonly regulated by *TP53* in either untreated or mafosfamide-treated condition. **B-D)** Enriched cellular component GO terms identified in the analysis of significantly regulated proteins by *TP53* expression in untreated (-) and mafosfamdie-treated (+) hMB cells. **B)** Sphere sizes represent the number of proteins associated to every GO term, while the color indicated the adjusted p-value (lower panel). **C, D)** Network interaction depicting the linkage of proteins and selected CC-GO terms identified in the comparison analysis between TP53-deficient and control hMB cells upon **C)** vehicle (untreated) or **D)** mafosfamide treatment. Sphere sizes represent the number of proteins associated to every GO term. Green and red nodes highlight down- and upregulated proteins respectively. Proteins involved in extracellular vesicle biogenesis or release have been highlighted.

### Extracellular Vesicle secretion from *TP53*-deficient lymphoma B-cells reduces the anti-tumor effector function of macrophages

Extracellular vesicles (EVs) have emerged as an important mechanism of cellular communication between tumor and stroma cells (Pegtel and Gould, 2019; Théry *et al*., 2018). To assess the relevance of the proteomic findings, we sought to characterize EVs produced by control and *TP53*-deficient lymphoma B-cells. EVs were isolated from lymphoma B-cell supernatants. The presence and size of the EVs was confirmed by transmission electron microscopy (Figure 5A) after enrichment and purification, showing vesicles in typical cup-shaped morphology with a size ranging from 50 to 200 nm. Loss of *TP53* did not alter vesicle size. However, nanoparticle tracking analysis (NTA) showed that *TP53*-deficient cells released significantly higher numbers of EVs than the control cells (Figure 5B). Then, following indications of the International Society for Extracellular Vesicles (ISEV) (Théry *et al*., 2018), we characterized EV-protein composition by immunoblot, detecting an enriched expression of the EV markers CD81, CD9, CD63 and Syntenin, compared to the respective cell source, while calnexin, a negative marker for EVs, was only presented on the cell lysate samples (Figure 5C). To determine whether *TP53*-deficient cell-derived EVs could affect macrophage properties, we exposed different types of macrophages to EVs isolated from *shCTRL* (*shCTRL*-EVs) or *shTP53* lymphoma B-cells (*shTP53-*EVs). First, we analyzed the EV-uptake by macrophages (Figure 5D). To this end, EVs labeled with the membrane DiD dye were added to J774A.1 or peritoneal macrophages. Our results showed that EVs derived from *TP53*-deficient cells were engulfed by macrophages in a similar way as the EVs released by the control cells (Figure S5A). Assessing the functional impact of EVs on ADCP, we observed that macrophages exposed to *shTP53*-EVs during the co-culture exhibited a significantly reduced phagocytic capacity of lymphoma B-cells, while they were non-significantly affected by *shCTRL*-EVs (Figure 5E). TP53-dependent inhibition on the ADCP persisted even when the concentration of *shTP53*-EVs added in the co-culture was reduced by 90% (Figure S5B). In parallel, we excluded EV-mediated effects on lymphoma B-cell viability (Figure S5C). To clarify if *shTP53*-EVs were affecting general macrophage phagocytic capacity, we performed a bead-based phagocytosis assay where macrophages were exposed to *shCTRL*-or *shTP53*-EVs 16 h prior to the adding of DyLight680 labeled latex beads (Figure S5D). Bead phagocytosis was unaffected by the presence of *shTP53*-EVs (Figure S5D). Thus, lymphoma B-cell EVs derived from *TP53*-deficient cells specifically abrogated Fc-receptor dependent macrophage anti-tumor effects, such as ADCP. Next,, to address the functional role of EVs on the lymphoma B-cell-macrophage crosstalk, we generated lymphoma B-cells unable to release EVs by depleting *RAB27A* expression via the CRISPR/Cas9 approach (Figure 5F)(Catalano and O’Driscoll, 2020). By NTA and protein concentration determination, we proved that *RAB27A* knockout cells (*RAB27A*-KO) showed deeply reduced ability to produce EVs (Figure 5G). Notably, cell viability and proliferation were unaffected by Rab27a loss. We assessed ADCP of *TP53*-deficient lymphoma B-cells lacking EV-release (*shTP53/RAB27A*-KO) and showed that the ADCP of EV-deficient tumor cells was significantly higher, than the *shTP53/RAB27A*-WT controls (Figure 5H). To validate our results *in vivo*, we used NSG mice transplanted with *hTP53*/*RAB27A*-KO or *shTP53*/*RAB27A*-WT hMB cells and treated the mice with CIT or PBS as a control. We observed that mice transplanted with lymphoma B-cells unable to release EVs (*hTP53*/*RAB27A*-KO) displayed significantly longer overall survival after CIT (CA+*RAB27A*-KO median = 21 ± 0.48 days vs. CA+*RAB27A*-WT median = 17 ± 0.47 days, p≤<0.0001) (Figure 5I). To further support our hypothesis, we also observed a significant increase in total number of EVs from TCL1/wt-mice vs TCL1/*Tp53*^*fl/fl*^ mice (Figure 5J), whilst also observing an increase from *TP53*-mutant primary CLL cells vs. wildtype cells (Figure 5K). In conclusion, we proved that EV formation is increased by loss of *TP53* and leads to suppression of macrophage effector function towards malignant cells.

**Figure 5.**
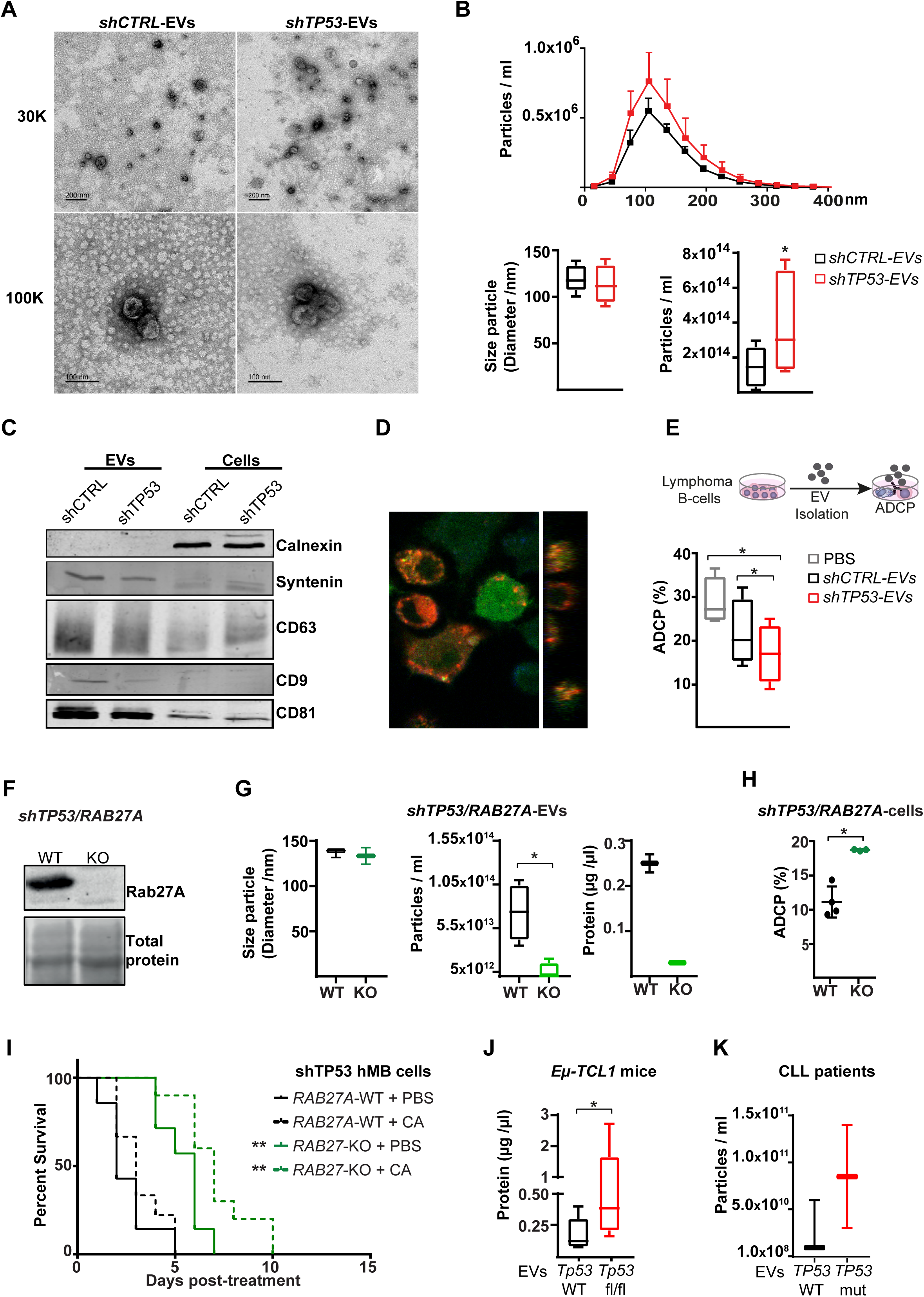
Characterization of features and functions of EVs derived from *TP53*-deficient and control lymphoma B-cells. **A-G)** EVs were isolated from *TP53*-deficient (*shTP53*-EVs) and control (*shCTRL-*EVs) hMB lymphoma B-cells. **A)** Representative TEM images.. 30k and 100k magnifications are shown. **B)** NTA EVs obtained by ZetaView. Distribution of the particles according to their size and concentration (upper panel). Box plots showing size particle and particle concentration of the isolated EVs (lower panel; n=6). **C)** Immunoblot analysis of the isolated EVs and their corresponding cell lysates using the indicated antibodies. **D)** Confocal image showing DiD-labeled EV (red) uptake by GFP^+^J774A.1 macrophages (green) after 16 h in culture. Orthogonal view of the same image confirms the intracellular presence of the EVs (right panel). **E)** Alemtuzumab-mediated ADCP of control hMB cells co-cultured with J774A.1 macrophages and *shCTRL*-EVs, *shTP53*-EVs or vehicle (PBS)(n=4). **F-I)** *RAB27A* expression was depleted by CRISPR/Cas9 approached in *shTP53* hMB cells. **F)** Immunoblotting detection of Rab27a protein in *shTP53/RAB27A*-WT and *-*KO hMB cells. The corresponding total protein staining of the samples as a protein loading control is shown. **G)** NTA of EVs obtained from *shTP53/RAB27A*-WT and *shTP53/RAB27A-*KO EVs hMB cells measured by ZetaView and BCA assay. **H)** Alemtuzumab-mediated ADCP of 3 different clones of both, *shTP53/RAB27A*-WT and *-*KO hMB cells co-cultured with J774A.1 macrophages. **I)** NSG mice were *i*.*v* injected with *shTP53/RAB27A*-WT or *-*KO hMB cells and *i*.*p* treated with a combination of cyclophosphamide and alemtuzumab (CA). PBS used as control treatment (n=7-10). **J)** Protein determination by BCA of EVs derived from CD19^+^cells isolated from the spleen of sick TCL1/wt *and* TCL1/*Tp53*^*fl/fl*^ mice (n=5-8). **K)** Concentration of EVs isolated from primary CLL patient cells of vs. TP53 mutant samples (n=2-3) (\**p*□<□0.05; \*\**p*□<□0.05).

### PD1/PDL1 signalling pathway inhibition restores the CIT-induced macrophage phagocytosis of *TP53*-deficient lymphoma B-cells

Given the diminished phagocytic capacity of macrophages in the presence of EVs produced by *TP53*-deficient malignant B-cells, we considered the possibility that the EVs secreted by *shCTRL* and *shTP53* cells displayed different expression levels of “don’t eat me” and related immune checkpoint molecules. Our data confirmed absence of CD47, but the surface presence of CD200 and PDL1on EVs derived from leukemic B cells by microsphere-based flow cytometry (Figure 6A). To assess whether PDL1 loaded on the EVs might be mediating the diminished macrophage phagocytic function, we blocked PDL1 expression on the EVs with the anti-PDL1 antibody atezolizumab before adding the EVs to the co-culture of macrophages and leukemic B-cells (Figure 6B). Remarkably, neutralizing PDL1 on the *shTP53*-EVs restored macrophage anti-tumor capacity assessed by ADCP. This observation was confirmed by EVs obtained from *PDL1*-knock-out (*PDL1*-KO) hMB cells, which do not impair macrophage anti-tumor function (Figure 6C, D). Thus, PDL1 expression on *TP53*-deficient lymphoma B-cell-derived EVs mediate suppression of macrophage phagocytic activity.

**Figure 6.**
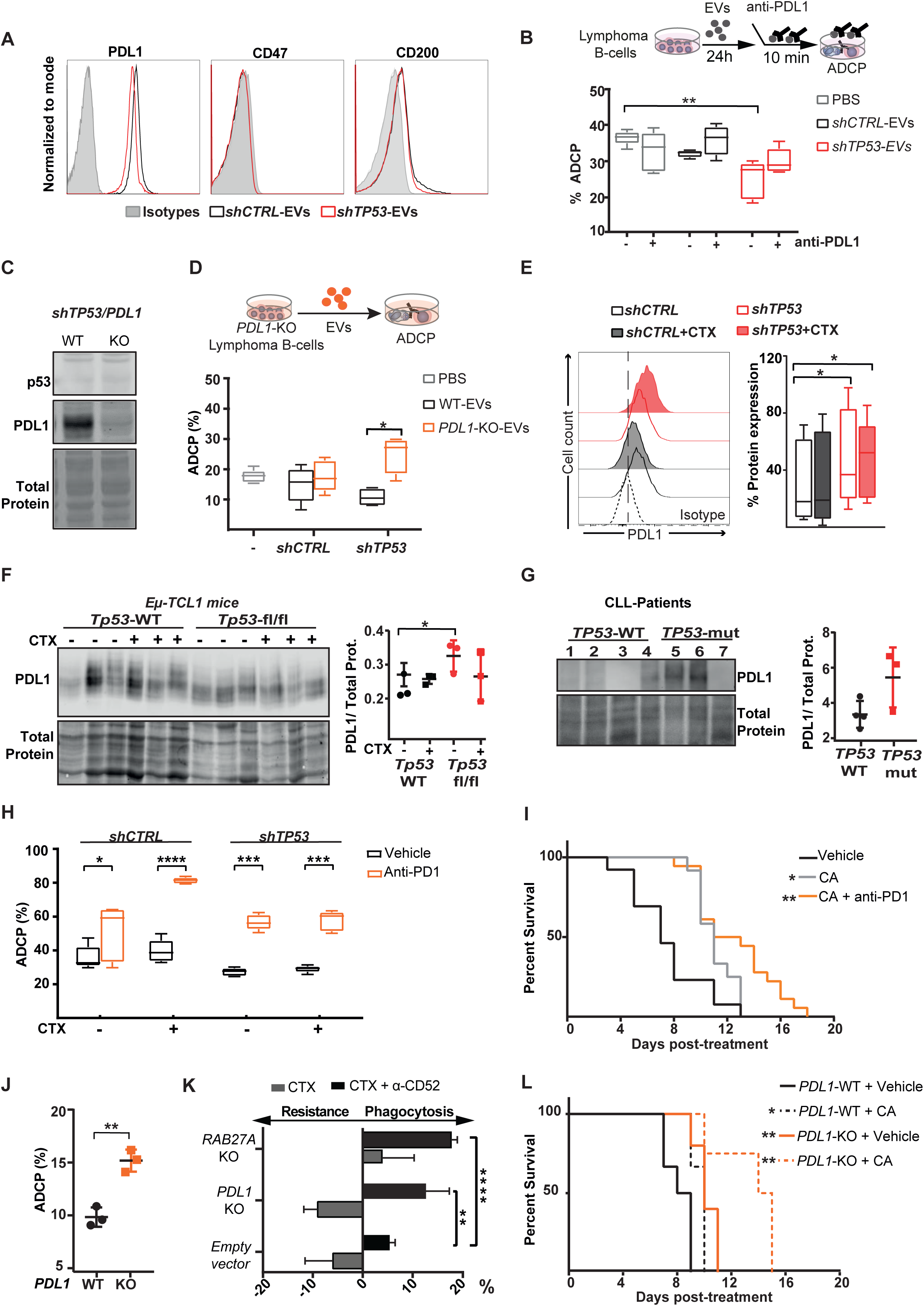
Identification of *shTP53*-EVs and PDL1 expression roles on the resistance to chemotherapy in *TP53*-deficeint B-cell lymphomas. **A)** Flow cytometer analysis of *shCTRL-* and *shTP53*-EVs bound to polystyrene microspheres and stained for the indicated proteins or the correspondent isotype control. **B)** Alemtuzumab-mediated ADCP of normal hMB cells and J774A.1 macrophages in the presence of EVs derived from *shCTRL* and *shTP53* hMB cells that were shortly pre-incubated (10 min) with anti-PDL1 antibody (atezoluzumab) prior to their addition to the co-culture. The same concentration of anti-PDL1 antibody diluted in PBS was used as control. Data shows 1 representative experiment with 5 replicates. **C)** Immunoblotting detection of p53 and PDL1 protein in *shTP53/PDL1*-WT and *-*KO hMB cells. The corresponding total protein staining of the samples as a protein loading control is shown. **D)** Alemtuzumab-mediated ADCP of normal hMB cells and J774A.1 macrophages in the presence of EVs derived from *shTP53/PDL1*-WT and -KO hMB cells. PBS was used as control. Data shows 1 representative experiment with 5 replicates. **E)** Flow cell cytometer detection of PDL1 in *shCTRL* and *shTP53* hMB cells, treated with mafosfamide (CTX) or vehicle for 24 h. Left panel shows half offset histogram representation of one representative experiment. Right panel displays PDL1 protein expression percentage from 6 independent experiments. **F-G)** PDL1 protein levels detected by Western blot and normalized to total amount of protein in each lane of **F)** CD19^+^ cells obtained from TCL1/wt *and* TCL1/*Tp53*^*fl/fl*^ leukemic mice treated with 10mg/kg cyclophosphamide (CTX) or PBS for 24 h (n=3) and, **G)** of *TP53*-WT (n=3) and *TP53*-mutated CLL samples (n=4). In both cases, densitometry analysis is shown (right panels). **H)** Anti-CD20 (18B12)-mediated ADCP of *shCTRL* and *shTP53* hMB cells pretreated or not with mafosfamide and co-cultured with peritoneal macrophages exposed to anti-PD1 antibody (GS-696882) from 4 h prior to the assay. Data shows 1 representative experiment with 5 replicates. **I)** Survival curve of NSG mice *i*.*v* injected with *shTP53* hMB tumor cells and treated *i*.*p* with cyclophosphamide and alemtuzumab (CA) or CA and anti-PD1 antibody (GS-696882). PBS was used as control treatment. (n=12-18). **J)** Alemtuzumab-mediated ADCP of different *shTP53*/*PDL1*-WT and *-*KO clones co-culture with J774A.1 macrophages (n=3). **K)** Number of mafosfamide (CTX) pre-treated GFP+ hMB cells (*shTP53*/*PDL1-KO, shTP53*/*RAB27A-KO* and the corresponding *shTP53* empty vector) remaining in the co-culture after ADCP assay performed in combination or not with alemtuzumab and normalized to the amount of hMB cells in a hMB single culture. Data shows 1 representative experiment with 5 replicates. **L)** Survival curve of NSG mice *i*.*v* injected with *shTP53/PDL1-*WT or *shTP53/PDL1-*KO hMB tumor cells and *i*.*p* treated after 10 days with cyclophosphamide and alemtuzumab (CA) or PBS as control treatment. (n=3-6). (\**p*□<□0.05; \*\**p*□<□0.01; \*\*\**p*□<□0.001; \*\*\*\**p*□<□0.0001).

As the above-mentioned results pointed towards a chemotherapy-priming effect on *TP53*-deficient lymphoma B-cells by displaying a “don’t eat me” phenotype, we addressed the cell surface expression patterns of potential suppressors of macrophage phagocytosis in the context of alkylating treatment *in vitro* (Figure 6E and Figure S6A). While CD47 expression showed stable and unaltered expression in all conditions, we could identify a reduction of CD200 expression after mafosfamide treatment in both, *shCTRL* and *shTP53* hMB cells. Interestingly, our results showed that *shTP53* hMB cells exhibited a significantly higher PDL1 protein expression than the control cells, and that this PDL1 upregulation is maintained after chemotherapy exposure (Figure 6E). We could corroborate that primary *TP53*-deficient malignant B-cells also expressed higher PDL1 protein levels than the respective controls, shown by analyzing both samples from TCL1-mice (Figure 6F) and from CLL patients (Figure 6G). This observation prompted us to evaluate the relevance of PD1/PDL1 axis in the resistance to chemo-immunotherapy. We performed a phagocytosis assay in the presence of anti-PD1 antibody in combination or not with chemotherapy (Figure 6H). The addition of anti-PD1 antibody to the co-culture significantly improved the ADCP of both *shCTRL* and *shTP53* hMB cells. It is noteworthy that the triple combination yielded phagocytosis of the control cells of more than 80% in the *in vitro* ADCP assay (Figure 6H). This finding was confirmed by using the *Myd88* p.L252P-driven DLBCL model, where PD1/PDL1 inhibition restored the resistance to chemotherapy observed on the ADCP of *TP53*-deficient tumor cells (Figure S6B). In addition, the use of the triple therapy approach (anti-PD1 antibody, cyclophosphamide and alemtuzumab) in the humanized mouse model of DHL with *TP53*-deficient lymphoma cells, revealed a significantly increased overall survival of the mice, compared to cyclophosphamide/alemtuzumab combination therapy (CA median = 11 ± 0.54 days vs. CA + anti-PD1 median = 13 ± 1.40 days, p = 0.023) (Figure 6I). Considering the lack of B and T cells in the immune deficient NSG-mice used in our model, the improvement on therapy response upon anti-PD1 treatment also demonstrated that macrophage tumor support depends on PD1/PDL1 activation. To clarify the PD1/PDL1 importance in the abrogation of macrophage anti-tumor functions on *TP53*-deficient lymphomas, we analyzed the ADCP of *shTP53/PDL1*-KO hMB cells (Figure 6J, K). We demonstrated that *PDL1* deletion favored the ADCP of *TP53*-defcient cells *in vitro* (Figure 6J), showing a higher phagocytic response of macrophages upon CIT (Figure 6K). Finally, we evaluated the role of PDL1 in the resistance to chemotherapy *in vivo* by injecting *shTP53/PDL1*-KO hMB cells into NSG mice (Figure 6L). Our results showed a significantly prolonged survival of *shTP53/PDL1*-KO lymphoma *vs. shTP53*-hMB lymphoma bearing mice (PBS+*PDL1*-KO median = 10 ± 0.55 days *vs*. PBS+*PDL1*-WT median = 8 ± 0.61 days, p = 0.007). Importantly, CIT response was highly effective in mice injected with *shTP53*/*PDL1*-KO tumor cells (Figure 6L) (PBS+*PDL1*-WT median = 10 ± 0.58 vs. CA+*PDL1*-KO median = 14 ± 1.67 days, p = 0.005). Thus, CIT treatment in combination with PD1/PDL1 inhibition improved *TP53*-induced therapy resistance. In conclusion, the upregulation of PDL1 on *TP53*-deficient tumor cells appears to be a resistance mechanism amenable for therapeutic targeting in B-cell malignancies in addition to CIT.

## Discussion

The concept of p53 as the central node in the DDR and “guardian of the genome” has recently experienced an expanded perception towards cell-non-autonomous effects in relation to the interaction with the TME (Levine, 2020; Ou *et al*., 2019). The functional impact of p53 on the TME in several tumor models and its functional restoration could reinstate immune surveillance in liver cancer (Raj and Attardi, 2013; Xue *et al*., 2007). Here, we identify *TP53* expression in B-cell lymphoma cells as a functionally suppressive switch in the interaction with macrophages of the TME.

In previous work, we could determine the essential role of cytokine release from lymphoma cells upon alkylating chemotherapy for sensitization of macrophage effector functions as the synergistic mechanism in CIT (Lossos *et al*., 2019; Pallasch *et al*., 2014b). Here, we show that an intact DDR is required for sensitization of this mechanism of synergy. We demonstrate that disruption of *TP53* function as a central node of the DDR leads to resistance against therapeutic antibodies in the context of CIT. Corresponding to this observation, loss or mutation of *TP53* is the most important resistance mechanism to CIT in CLL (Campo *et al*., 2018; Eichhorst *et al*., 2016; Hallek *et al*., 2010). Moreover, the regulatory effect of supernatants from *TP53*-deficient lymphoma cells on macrophage effector function was caused by a profoundly elevated secretion of EVs and an increase of the immune checkpoint PDL1 expression. In particular, we identify the axis of *TP53* dependent regulation of extracellular vesicle release and regulation of PD-L1 expression.

Multiple mechanisms of controlling PDL1 expression have been described in malignant cells. Likewise, *TP53*-dependent control of PDL1 expression has been previously described via miRNA-34a regulation of the PDL1 3’UTR in small-cell lung cancer cells (Cha *et al*., 2019; Cortez *et al*., 2016). *TP53* status has furthermore been hypothesized to predict response to checkpoint inhibitor therapy of malignant disease (Assoun *et al*., 2019; Dong *et al*., 2017). The functional relevance of immune oncology drugs targeting the PD-1/PDL1 axis have so far mainly been attributed to T-cell activation and adaptive immunity (Sanmamed and Chen, 2018; Topalian *et al*., 2015). However, there is similar evidence that the PD1/PDL1 checkpoint regulates macrophages in the tumor microenvironment (Gordon *et al*., 2017). We have now identified PDL1 to inhibit macrophages to effectively engulf lymphoma cells, suggesting PDL1 as a novel resistance factor to CIT.

In this line, expression of the immune checkpoint PDL1 was described as an integral component parameter in a biomarker score indicating poor response to immunochemotherapy of lymphoma (Keane *et al*., 2015). In DLBCL, patients with high expression of PD-1/PDL1 on T cells and macrophages had significantly poorer survival after R-CHOP (Xu-Monette *et al*., 2019). In this context, and in line with our data, it has been recently shown that loss of *TP53* increases PD-L1 in a murine lymphoma model of DLBCL, inducing immune evasion, which could be overcome with PD1-blockade (Pascual *et al*., 2019). Importantly, our data now indicates that PDL1 dependent immune suppression is independent of direct cell-cell interactions. We observe the increase of released EVs exposing PDL1 to mediate the suppression of phagocytic function of effector macrophages. Notably, we see this novel phenomenon both in DLBCL and CLL murine models with loss of *TP53* function as well as in CLL patient samples. Using genetic targeting of PDL1 on EVs, applying EV-deficient cells and immune checkpoint inhibitor blockade of EVs; we provide direct evidence of the decisive functional role of PDL1-positive EVs in regulating macrophage effector function and treatment response to CIT.

TP53 has been described to regulate EV secretion (Yu *et al*., 2006). More specifically it has been proposed that mutant *TP53* colon cancer cells reprogram macrophages to tumor supporting macrophages via miR-1246 delivered by EVs (Cooks et al., 2018). Colorectal cancer–derived EVs were described to regulate polarization of tumor-associated macrophages by the miR-145 which is transcriptionally induced by p53 (Sachdeva *et al*., 2009; Shinohara *et al*., 2017). As for the context of chemotherapy and DNA damage, EV-based metabolic reprogramming by tissue-infiltrating macrophages underlines the role of EVs in cellular crosstalk (Goulielmaki *et al*., 2020). Also, an increase of EV-shedding was demonstrated to promote melanoma growth after chemotherapy (Andrade *et al*., 2019). Melanoma derived exosomes were identified to carry the immune checkpoint ligand PDL1 and thereby contribute to immunosuppression (Chen *et al*., 2018). So far, no specific therapeutic addressing EV-mediated effects has been identified, nevertheless immune checkpoint inhibitors are offering the opportunity to target the EV-mediated checkpoint control. Proving our hypothesis, we were effectively improving CIT by targeting the PDL1 immune checkpoint and significantly enhancing ADCP macrophage effector function.

Future directions form this work point toward clinical evaluation of CIT combination with checkpoint inhibitors in order to overcome PDL1-associated resistance. Previous trials have shown that combination of chemotherapy with immune checkpoint inhibitors is clinically safe and showed 77% complete response rate in a recently published phase II trial combining pembrolizumab with R-CHOP (Smith *et al*., 2020). The loss of p53 function could serve as a future biomarker for intensified treatment and should be evaluated as a predictive marker in prospective clinical trials. Finally, EV-mediated resistance mechanisms in cancer therapy are amenable for specific targeting by systemic targeted therapies.

## Material and Methods

### Cell lines and primary patient material

The study was approved by the ethical commission of the medical faculty of the University of Cologne (reference no. 13-091) and an informed written consent was obtained from all patients. Primary CLL patient samples were isolated from peripheral blood as previously published (Frenzel et al., 2011; Herling et al., 2018). Primary CD14-positive monocytes were obtained from healthy donor buffy coats (Blood bank Cologne, Germany) and differentiated to macrophages. For this, human recombinant M-CSF (Miltenyi, Berg. Gladbach, Germany) was added (final concentration of 50U/ml) and cells were incubated for 3 days. Then, 40% of the original medium volume was added with the respective amount of M-CSF and cells were incubated for 3 days before they were used for functional experiments.. To isolate peritoneal macrophages, 8 to 24 weeks old wildtype C57BL/6 mice (Charles River, USA) were injected *i*.*p* with thioglycolate and macrophages were obtained via peritoneal lavage after four days. In parallel, bone marrow derived macrophages (BMDM) were obtained from the same mice as explained before (Pallasch et al., 2014a). Peritoneal macrophages, BMDM and the murine macrophage cell line J774A.1 were cultured in DMEM (Gibco) supplemented with 10 % FBS (Biochrom GmbH) and 1% Pen/Strep (Gibco). BMDM media was supplemented with 15% Feeder media 1 (supernatant derived from confluent unstressed L929 fibroblasts) and 15% Feeder media 2 (supernatant derived from confluent stressed L929 fibroblasts). The human-MYC/BCL2 (hMB) cell line (strain 102), generated by Leskov *et al*.(Leskov et al., 2013) was cultured in B-cell culture medium (BCM) composed of a 1:1 ratio of Iscove’s Modified Dulbecco’s Medium (IMDM) and DMEM, supplemented with 10% FBS, 1% P/S, 1% GlutaMAX and 1% β-Mercaptoethan. Myd88p.L252P-driven ABC-DLBCL (Myd88p.L252P) cells were obtained as described before (Knittel et al., 2016) and culture in BCM. HEK 293T derived amphotrophic phoenix and ecotropic phoenix cells were cultured in DMEM, supplemented with 10% FBS and 1% P/S.

### Isolation of myeloma-associated macrophages and myeloma cells from human samples

Bone marrow (BM) cells from patients with MM were stained with CD163, CD15 and CD138 antibodies. Macrophages (CD163+/CD15-) and myeloma cells (CD138+) were isolated by flow cytometry. Purity was greater than 95%. Each Patient gave informed consent prior to surgery or bone marrow biopsy, and the institutional ethics committee approved the study (Erlangen: Ref. number: 36_12 B, 219_14B, 200_12B).

### Generation of shRNA mediated knock downs in hMB cells

shRNA sequences were designed by using the gene specific NM number (CoreNucleotide; Pubmed database) in the siRNA Scales tool (http://gesteland.genetics.utah.edu/siRNA_scale) and the shRNA converting tool (http://katahdin.cshl.org:9331/homepage/siRNA/RNAi.cgi?type=shRNA#oligos=1). The corresponding direct and reverse oligos for every shRNA (Invitrogen; Thermo Fisher Scientific, Waltham, USA) were annealed and subcloned on a MLP plasmid vector which contains a green fluorescent protein sequence (GFP) (Jiang et al., 2011). Then, shRNAs were cloned into MLP plasmid backbone and the final shRNA-MLP plasmids were used to generate the different hMB DDR-knockdowns by retroviral transduction. Here, HEK 293T derived amphotrophic phoenix cells were transfected by calcium phosphate method with 25μM chloroquine and the corresponding shRNA-targeting MLP vector in combination with psPax2 (Addgene, #12260) and pMD2.G (Addgene #12259) plasmids, both gifts of Didier Trono. After 48h of incubation the virus containing medium was harvested, filtered through a 0.45μm syringe filter and used by spin infection (400g 32°C 45 min) of the hMB cells. Then, cells were incubated polybrene for 48h at 37°C before antibiotic selection undertaken with 3μg/ml puromycin (Invivo Gen, San Diego, CA, USA) for at least 1 week.

### Generation of shRNA mediated knockdowns in Myd88p.L252P cells

shRNAs (gift from MT Hermann; Koch Institute for Integrative Cancer Research, MIT, USA) (Jiang et al., 2011) were cloned into a MLP plasmid backbone. In a similar way than hMB cells, Myd88p.L252P cells were transduced with shRNA-MLP retroviral particles, obtained by calcium phosphate co-transfection of ecotropic phoenix cells with the corresponding shRNA-targeting MLP vector and the plasmids psPax2 and pMD2.G. Myd88p.L252P transduction was performed as explained above. Then, cells were incubated for 48h at 37°C before antibiotic selection.

### Generation of CRISPR mediated knockout constructs

For the generation of both, lentiviral-based human *PDL1* and *RAB27A* via CRISPR/Cas9, single-guide RNAs (sgRNA) specifically recognizing *PDL1* and *RAB27A* genes were selected respectively, using the web-based tool CHOPCHOP v3 (Labun et al., 2019). sgRNA oligonucleotides (Merck KGaA, Darmstadt, Germany) were cloned into the lentiCRISPR v2-Blast plasmid (LV-CRISPRblast) created by Mohan Babu (Addgene, #83480).

### Generation of stable hMB knockout cells by lentivirus infection

The corresponding LV-CRISPRblast plasmid (sgRNA-*PDL1* or sgRNA-*RAB27A)* was packaged into lentiviral particles using HEK 293T cells co-transfected with the plasmids psPAX2 and pMD2.G. Six hours after transfection, the medium was changed and subsequently collected at 24□h intervals till 72 h. *shCTRL* and *shTP53* hMB cells were transduced with 0.45 µm filtered lentivirus by spin infection (800g 32°C 2h). After infection for 48□h, the medium was replaced with fresh medium, and infected cells were selected with blastomycin (Invivo Gen). After a week, limited dilution in 96 wells was used to grow single cell clones.

### Western blot analysis

Whole cell and EV pellets were lysed with RIPA buffer (Cell Signalling Technology Cat.) containing phosphatase Inhibitor cocktail (100x, Thermo Fischer Scientific) and protease inhibitor cocktail, EDTA free (100x, Thermo Fischer Scientific). In the case of EV blots, EVs and matching cell pellets were also sonicated. Then, 10 µg of total protein of each sample was separated on 10% SDS-PAGE under reducing or non-reduction conditions, according to antibody manufacturer’s instructions, and transferred onto a nitrocellulose membrane (GE Healthcare, Freiburg, Germany). The membranes were blocked 1 h at RT (according to manufactures’ protocol) and incubated with the corresponding primary antibody overnight at 4°C. After washing, membranes were stained with secondary fluorescent dye-labeled antibodies (LI-COR Biotech., Bad Homburg, Germany) for 1h RT. Protein bands were detected at 700 or 800nm using the LI-COR Odyssey infrared imaging system. Protein loading was either normalized using antibodies against ß-actin and GAPDH or to total protein content detected by Ponceau acid red, or Revert(tm) 700 Total Protein Stain (LI-COR) according to manufacture instructions. Densitometry was performed with Image Studio Lite Ver 5.2 software.

### Antibody dependent phagocytosis assay (ADCP)

Macrophages were plated out in 96-well plates at an optimized concentration varying between the different cell types (1×10^5^ cells/ml for J774.A1 and 5×10^5^ cells/ml for peritoneal macrophages and human macrophages) and left them to attach for 6-16 h. Then, 100 µl of media containing lymphoma B-cells (1.5×10^5^ cells/ml for hMB and TCL1-derived cells and 5×10^6^ cells/ml for human CLL cells) were added to both, wells containing macrophages and wells without macrophages. Subsequently, cells were culture for 16 h in combination or not with the corresponding monoclonal antibody. To stimulate the ADCP of human cells the human specific monoclonal antibody alemtuzumab (Genzyme, Cambridge, MA, USA) was used at 10μg/ml. For murine lymphoma B-cells we used the specific murine anti-CD20 monoclonal antibody 18B12 (Biogen, Cambridge, MA; USA) at 20μg/ml or 50µg/ml. Each condition was performed with five replicates. For determination of ADCP, remaining GFP^+^ target cells (hMB or Myd88p.L252P) were analyzed using a MACSQuant VYB flow cytometer (Milentyi Biotech, Berg. Gladbach, Germany). Primary leukemic cells as human CLL cells or TCL1 cells were stained with CD19 fluorescent antibody specific for the respective species for 15min at 4°C before measurement by flow cytometry. The percentage of ADCP was calculated as follows: 100 - (100 x (cells/µL treated / cells/µL untreated)).

### Generation of conditioned media

For the generation of conditioned media tumor B-cells were incubated with 3 µM of mafosfamide sodium salt (Santa Cruz, Santa Cruz, CA, USA), 10 µM Nutlin-3A (Tocris Bioscience, Bristol, UK) or the corresponding amount of DMSO as vehicle control for 24 h. This mafosfamide concentration induced around 20% toxicity. Afterwards, cells were washed and plated out at a concentration of 4×10^6^ cells/ml for 24h. The generated supernatant was collected and centrifuged for 10 min at 2000 rpm and stored at −80°C for different experiments.

### ADCP assay for primary myeloma cells

Myeloma cells were labeled with CPD and co-cultured with macrophages in different effector target ratios (1:1, 5:1 and 10:1) in sterile FACS tubes in the presence or absence of Daratumumab (10µg/ml) for 24 hour (CM+10%FCS, 37°C, 5%CO2). An irrelevant IgG1 antibody was used as an isotype control. To distinguish between phagocytosed CPD positive MM cells and free MM cells macrophages were counterstained with anti-CD11b-FITC antibody. Absolute numbers of surviving MM cells were determined by flow cytometric analysis of CD11b-/CPD+ cells via 123count eBeads (ebioscience).

### Eµ-TCL1 leukemic mice

*Eµ-TCL1*/*Cd19*-*Tp53* wildtype (TCL1/wt) and, *Eµ-TCL1/Cd19* ^*Cre/wt*^*-Tp53* ^*fl/fl*^ (TCL1/*Tp53*^*fl/fl*^) were generated as described elsewhere (Knittel et al., 2017). Briefly, B cell-specific loss of *Tp53* in *Eµ-TCL1* mice was modeled by crossing the *Eµ-TCL1* allele with *Cd19*^*Cre*^ and *LoxP*-flanked *Tp53* allele on a mixed C57BL/6J-C57BL/6N background. Mice were treated *in vivo* with cyclophosphamide (10mg/kg) or PBS when reaching 8-12 weeks old and 30% of leukemic cells (CD45^+^CD19^+^CD5^+^ cells) in the peripheral blood. After 24h, mice were sacrificed and the spleen leukemic cells were obtained for *ex vivo* and *in vitro* experiments. Animal experiments were conducted with permission of the Landsamt für Natur, Umwelt und Verbraucherschutz Northrhine-Westphalia under the file numbers 84-02.04.2014.A146, 81-02.04.2019.A009 and Uniköln_Anzeige§4.16.009.

### Humanized DHL mouse model

For *in vivo* experiments 8-14 weeks old male and female NOD.Cg-*Prkdc*^scid^ *II2rg*^tm1Wjl^/SzJ (NSG, Jackson Laboratory, USA) immunodeficient mice were injected *i*.*v*. with 1×10^6^ hMB cells diluted in 100 µl PBS. Unless otherwise stated, 10 days after injection mice were treated *i*.*p*. on three consecutive days with alemtuzumab (day 10: 1 mg/kg, day 11 and day 12: 5 mg/kg) and one dose of cyclophosphamide (day 11: 100 mg/kg or PBS as control). For the treatment with anti-PD1 antibody GS-696882 (Gilead Sciences; USA), mice received an *i*.*p* injections of anti-PD1 (10 mg/kg) on day 10, 12 and 14 of the experiment. Disease progression was monitored by weekly blood sampling and daily scoring of the mice. Spleen, bone marrow and ovaries were harvested and dissociated by cell strainers with PBS and lysed with 5 ml ACK lysis buffer for 3 min at RT. Then, the concentration of GFP^+^ hMB cells were detected by flow cytometry. Animal experiments were conducted with permission of the Landsamt für Natur, Umwelt und Verbraucherschutz Northrhine-Westphalia under the file numbers 84-02.04.2016.A119 and Uniköln_Anzeige§4.16.009.

### Multiple cytokine array (Eve Technology)

100μl conditioned media from *in vitro* mafosfamide and vehicle control treated *shCTRL* and *shTP53* hMB cells were sent to Eve technology (Calgary, Canada) on dry ice for bead based multiple cytokine array. Samples were measured in duplicates, concentrations is displayed in pg/ml.

### Single cell RNA-Seq – Sample preparation

Splenocytes from all mice were depleted of CD19^+^ B-cells using the CD19 MicroBeads (Miltenyi Biotech). CD19^-^ splenocytes from all mice were checked for viability via Trypan Blue using a Neubauer chamber. If the viability was <70%, the cells were negatively sorted for viable cells using the Miltenyi Dead Cell Removal kit (Miltenyi Biotech). The cell concentration for each sample was adjusted to allow for a target output of ∼10000 cells/sample. Following which, single cells from all samples using the Chromium Controller were encapsulated into emulsion droplets (10x Genomics). The Chromium Single Cell 3’ v2 Reagent Kit was used to construct the scRNA-Seq libraries, according to the manufacturer’s protocol. Libraries were sequenced using the NovoSeq 6000 (Illumina; San Diego, CA; USA).

### Single cell RNA-Seq – Data analysis

Samples were analyzed with the M3K in-house processing pipeline. For each sample, barcodes and UMIs were extracted from sequences in file1 and concatenated to corresponding sequence names in file2. Barcodes having a mismatch at a single nucleotide position compared to the reference list of barcodes provided by 10x Genomics were corrected, where possible. Next, sequences from file2 were mapped to altered mouse genome (mm10) using STAR Aligner (version 2.5.2b) only keeping uniquely mapped reads (Dobin et al., 2013). Duplicate reads were identified as sequences within the same cell with the same UMI and same or slightly variable start position (between 1 and 3 nucleotides), as described in Sena et al. and were excluded from further analysis (Sena et al., 2018). In addition, UMIs at each exonic region with 1 nucleotide mismatch from a set of UMIs residing at that exon were considered as duplicate reads and were consequently removed. First and second derivatives in combination with curve smoothing were used to identify a point where the number of quantified reads per cell drastically drops marking it a cutoff for the number of viable cells. Resulting matrix containing barcodes (cells) and genes were then used as input for further downstream analysis using Seurat R package (Butler et al., 2018; Stuart et al., 2018). Initially, all samples underwent pre-processing, removing cells with low (<200) and high (>2000-3000, dependent upon sample) feature counts, as well as removing cells with >5% mitochondrial gene counts. All samples were log normalized prior to highly variable feature identification, where the top 2000 variable features were taken forward for data scaling and linear dimension reduction. After which, the JackStraw procedure was implemented to determine the number of principle components to be taken forward for clustering. We leveraged the graph-based clustering approach utilized in the Seurat v3 package, followed by non-linear dimension reduction using Uniform Manifold Approximation and Projection (UMAP) for each sample, followed by differential gene expression analysis. For sample integration, we utilized the Seurat v3 Integration strategy published by Stuart *et al*. (2019). Briefly, we conducted sample pre-processing, which entailed log-normalization and variable feature identification across all samples, followed by the identification of “anchors”, using default parameters. Following which, we integrated the samples using these anchors, creating a “batch-corrected” expression matrix to conduct clustering, differential gene expression analysis to identify cell types (using the integrated assay), and downstream analysis of the treatment groups within clusters (using the RNA assay). Downstream analysis of individual clusters and differential expression between treatment groups was conducted within Seurat, by subsetting each representative cluster, renaming the identity to match the treatment group, and repeating the FindAllMarkers step from the standard protocol. Here, Gene Ontology (GO) annotations were assessed using the Gorilla online tool (http://cbl-gorilla.cs.technion.ac.il/) (Eden et al., 2009) Briefly, the list of significant up-/downregulated genes from each test were uploaded to the tool, using the full background gene list produced from the relevant test. Pathways were considered significantly up-/downregulated with *P*<0.05.

### Label-free LC-MS sample preparation and analysis

For the label-free LC-MS approach, 10×10^6^ *shCTRL* and *shTP53* hMB cells were plated out in serum-freem media and treated with 3 µM mafosfamide or vehicle (DMSO) for 6 h. Then, cells were centrifuged 5 min 1350 rpm and cell pellet was washed with cold PBS through 2 centrifugations rounds 5 min at 1500 rpm. Cell pellet were lysate with RIPA buffer (Cell Signalling Technology Cat.) containing phosphatase inhibitor cocktail and protease inhibitor cocktail (Thermo Fischer Scientific), followed by sonication with bioruptor (Diagenode) for 10 min at 4°C lysates cleared by centrifugation. 25 µg protein was precipitated with ice-cold acetone overnight at −20°C. Samples were then resuspended in 6 M Urea/ 2 M Thiourea, reduced with DTT and carbamidomethylated with IAA. LysC (Wako) and trypsin (Promega) were added for overnight digestion.

### LC-MS/MS Analysis

Proteomic analysis was performed using an Easy nLC 1000 ultra-high performance liquid chromatography (UHPLC) coupled to a QExactive plus mass spectrometer (Thermo Fisher Scientific) with the previously described settings (Pla□Martín et al., 2020). The raw files were processed using MaxQuant software (v1.5.3.8) and its implemented Andromeda search engine (Cox et al., 2011). The mass spectrometry proteomics data have been deposited to the ProteomeXchange Consortium via the PRIDE [1] partner repository with the dataset identifier PXD019687.

To establish a reproducible and transparent protocol, an in-house pipeline integrating several R packages (DEP (Zhang et al., 2018), limma (Ritchie et al., 2015), clusterProfiler (Yu et al., 2012), ggplot2 and GGally (Wickham, 2009)) was written and followed. The pipeline includes pre-processing steps as filtering of potential contaminants, removal of proteins with too many missing values (only proteins with intensities quantified in 3 replicates of at least one condition were kept). Normalization was performed using variance stabilizing transformation (vsn) and remaining missing values were classified in two categories as described in the MSnbase R package(Gatto and Lilley, 2012). The ones resulting from the absence of detection of a feature, despite being present at detectable concentrations were handled as missing at random (MAR) and imputed with maximum likelihood-based method (MLE) using the expectation-maximization algorithm. However, biologically relevant missing values resulting from the absence of low abundant ions (below the instrument detection limit) were classified as missing not at random (MNAR) and imputed with a left-censored approach using a deterministic minimal value (MinDet). Information regarding whether a protein normalized intensity value was imputed or not can be found in Supplementary Table 2.

Subsequently, differentially expressed proteins were identified using limma R package using a p-adjust ≤ 0.05 and 1 ≤ log_2_ fold change ≤ 1. Afterwards, GO term and KEGG pathway enrichment analyses of differentially expressed proteins were performed using clusterProfiler R package with a p-adjust ≤ 0.1. Visualization of results was conducted using ggplot2 and GGally R packges.

### EV isolation

EVs were extracted from supernatant of cells cultured 24 h at a concentration of 16.6 x10^6^ cells/ml in serum-free CD293 media supplemented with Glutamax 1%. Cell viability (85-100%) was confirmed after the 24 h incubation. Supernatants were subjected to a serial of centrifugation steps (1200 rpm 5 min, 2900 rpm 10 min and 3500 20 min) and passed through a 0.22 µm PES membrane filter (VWR). To enrich EV concentration we used the Total Exosome Isolation kit (Thermo Fisher Scientific) according to the manufacturer’s instructions. In brief, supernatants were incubated over night with 0.5 volumes of Total Exosome Isolation reagent. After incubation, samples were at 10000 g for 1 hour at 4°C followed by pellet resuspension in 1.5 ml of cold PBS (1X). Finally, EVs were ultracentrifuged (Type 45 Ti rotor, k-Factor 133, Beckman Coulter) with 100000 g for 90 min 4°C. Pellets were resuspended in cold PBS accordingly to the initial supernatant volume (1 µl PBS for 1 ml supernatant) and dissolved by pipetting with a 1 ml syringe (Henke-Sass Wolf, Germany) with 26G 7/8 inch needle (Terumo, Germany). EVs were immediately used for ADCPs assays or stored at −80°C. For Western blot analysis, EV pellet was directly lysed.

### Transmission electron microscopy (TEM) imaging of EVs

Formvar-coated copper grids (Science Services, München) were loaded with 5 µl of diluted sample (1:5) containing 0.5 µg protein as determined by BCA protein assay. The grids and samples were incubated for 20 min before being fixed with 2% paraformaldehyde for 5 min. Samples were washed with PBS and fixed again with 1% glutaraldehyde for 5 min, washed with Milli-Q water and contrasted for 4 min with 1.5% uranyl acetate. Images were acquired using a Gatan OneView 4K camera mounted on a Jem-2100Plus (Jeol) operating at 200kV.

### Nanoparticle tracking analysis (NTA)

The concentrations and size distribution of EVs were analyzed by NTA using the ZetaView instrument (Particle Metrix, Germany). All samples were diluted in PBS to a final volume of 1 ml. Ideal measurement concentrations were found by pre-testing particle per frame value (100–300 particles/frame). We used the manufacturer’s default software settings for EVs and proceeded as explained before (Bachurski et al., 2019).

### Macrophage uptake of DiD-labelled EVs

EVs derived from wildtype hMB cells were labeled (1:100) with Vybrant(tm) DiD Cell-Labeling Solution (Invitrogen) 20 min 37°C, followed by a wash step with PBS and 110000 g ultracentrifugation (90 min, 4°C). EV pellet was resuspended in PBS as explained above. To evaluate the uptake of EVs by confocal imaging, 2.5×10^5^ GFP^+^J774A.1 macrophages were overnight plated out. DiD-EVs were added to the cell cultures for 0, 1, 2, 4, 8, 16 and 24 h at 37°C. Then, cells were washed with PBS (1X), fixed with 4% PFA 10 min, washed and analysed by a Leica TCS SP8 microscope. To detect engulfed EVs by flow cytometry, 50 µl of DiD-EVs were added to 5×10^5^ cells/ml J774A.1 or peritoneal macrophages for 16 h 37°C. Cells were washed with PBS, carefully scraped and stained with BrilliantViolet421 conjugated anti-CD11b antibody (Biolegend). Macrophage uptake of DiD-EVs was measured using the MACSQuant X flow cytometer (Miltenyi Biotech.) and data were analyzed with FlowJo v10.6.2 software.

### Microsphere-based flow cytometry of EVs

EVs were incubated overnight (4°C) with polystyrene microspheres 4.5 µm (Polysciences INC, Warrington, PA). The microspheres were blocked with 2% BSA (v/w) in PBS for 1 h in a thermoshaker (25°C, 800 rpm). Next, the samples were incubated 4°C 15 min with fluorescently labelled antibodies or isotyope, followed by a wash step with FACS buffer (PBS+ 2% FBS+ 2mM EDTA) and centrifugation at 2000 rpm 3 min 4°C. Microspheres were analysed with MACSQuant X flow cytometer (Miltenyi Biotech.)

### Polybead Phagocytosis Assays

Polybead Amino Microsphere 3 μm latex beads (Polysciences INC) were labeled with the fluorescent dye DyLight680 mono-N-hydroxysuccinimide (NHS) ester-activated. The lyophilized powder was dissolved in 0.1 M sodium carbonate (pH 9.0) to a final concentration of 100 mg/ml. Five-hundred microliters of the latex beads were centrifuged at 3000 g for 5 min and the pellet was resuspended in 500 μl 0.1 M sodium carbonate (pH 9.0). Three-hundred micrograms of DyLight680 NHS ester were added and incubated for 2 h at room temperature while rotated. Afterwards, the labeled beads were centrifuged at 3000 g for 5 min and washed three times in 20 ml PBS (with the same centrifugation conditions). Finally, beads were resuspended in 1 ml PBS and correct labeling as well as the concentration of the beads was measured and calculated via flow cytometry.

### Statistics

Data were analyzed using GraphPad Prism 8.0 Prism software (San Diego, CA, USA) and SPSS. Results are expressed as mean ± SD and means were compared by Student’s t-test, Mann-Whitney U-test, and Wilcoxon tests as appropriate. Statistical comparison between groups was performed using one-way ANOVA multiple comparison test in the ADCP assays, or the Kruskal-Wallis test for non-gaussian distributed data. Kaplan Meier survival analysis was performed using pairwise Log-rank (Mantel-Cox) test. Differences were considered statistically significant at p-values less than 0.05 (*, p ≤ 0.05; **, p ≤ 0.01; ***, p ≤ 0.001; ****, p ≤ 0.001).

## Acknowledgements

We are indebted to our patients who contributed tissue and blood samples to this study. This work was supported by the German research foundation (DFG; KFO 286) RP5 CPP was supported by the ‘Foerderprogramm Nachwuchsforschungsgruppen NRW 2015–2021, CAP Program of the Center for Molecular Medicine Cologne and a research grant by Gilead Sciences, the German-Israeli Foundation for Research and Development (I-65-412.20-2016 to H.C.R.), the Deutsche Forschungsgemeinschaft (KFO-286-RP2 to H.C.R.), the Deutsche Jose Carreras Leukämie Stiftung (R12/08 to H.C.R.), the Else Kröner-Fresenius Stiftung (EKFS-2014-A06 to H.C.R., 2016_Kolleg.19 to H.C.R.), the Deutsche Krebshilfe (1117240 and 70113041 to H.C.R.) and the German Ministry of Education and Research (BMBF e:Med 01ZX1303A to H.C.R.). H.B. was supported by Wilhelm-Sander Foundation and by the German Research Foundation (DFG BR 4775/2-1). We are grateful for technical assistance from Yvonne Meyer, Sophie Neuber, support from Katrin Reiners, and the CECAD Imaging and animal facilities.

## Supplementary figure legends

**Supp.Figure S1. Validation of shRNA-targeted lymphoma B-cells. A, B)** Western blot detection of the corresponding DNA damage response proteins in **A)** shRNA-targed hMB cells and **B)** *Myd88p*.*L252P* cells. Beta-actin was used as protein loading control. **C)** Phagocytosis of *shCTRL* Myd88p.L252P cells by J774A.1 macrophages, peritoneal macrophages or bone marrow derived (BMD) macrophages. *Myd88p*.*L252P* cells were pre-treated or not with mafosfamide (CTX; 1.64 µM) and 50µg/ml of monoclonal anti-CD20 antibody (18B12) was added or not to the co-culture. After 16hrs, the remaining cells were enumerated by flow cytometry and normalized to the corresponding number of *Myd88p*.*L252P* cells present in single cell cultures. Data show one representative experiment of four independent experiments. (\**p*□<□0.05)

**Supp.Figure S2. *TP53*-inuduced expression on hMB cells with disrupted DDR signaling. A)** Determination of p53 activation levels in *shCTRL*and indicated DDR-knockdowns hMB cell lines after treatment or not with 3μM mafosfamide (CTX) for 24h. Expression is normalized to GAPDH as loading control. Right panel shows densitometry analysis of CTX-induced p53 activation fold change (mean ± SD) of four independent experiments. **B)** Western blot detection of p53 protein levels upon 10µM nutlin-3A or vehicle treatment on hMB-knockdowns for the mentioned DDR proteins. GAPDH was used as protein loading control. Densitometry analyses of nutlin-3A-induced p53 activation fold change (mean ± SD) of 3 independent experiments (right panel).

**Supp.Figure S3. Crosstalk alterations between *TP53*-defienct lymphoma B-cells and macrophages in the TME. A)** Heatmap representation of mafosfamide-induced cytokine expression on *shCTRL* and *shTP53* hMB cells obtained with a multiple cytokine array (n=2). **B)** Levels of VEGF, TNFα, IL8 and CCL4 produced by *shCTRL* and *shTP53* hMB cells treated or not with mafosfamide (n=2). **C-E)** TCL1/wt *and* TCL1/*Tp53*^*fl/fl*^ leukemic mice were *i*.*p* injected 10 mg/kg cyclophosphamide (CTX) or PBS for 24 h (n=1). TME CD19^-^cells were isolated from the spleens of each treatment group, followed by single-cell RNA sequencing (scRNA-seq). **C)** UMAPdimension reduction plot of every treatment group. **D)** Percentage of cell subtypes contained in the TME of the four conditions. **E)** Heatmap showing fold change gene expression induced by CTX of significant genes in each macrophage clusters (MΦ-C1, MF-C2 MΦ-C3 and MΦ-C4). (\**p*□<□0.05; \*\**p*□<□0.01; \*\*\**p*□<□0.001; \*\*\*\**p*□<□0.0001).

**Supp.Figure S4. Proteomic modifications induced by chemotherapy in *TP53*-deficient lymphoma B-cells. A-D)** *shCTRL* and *shTP53*hMB cells were treated with 3µM mafosfamide or vehicle (untreated) for 6 h and their proteomic profile were analyzed by mass spectrometry label-free protein quantification (n=3). **A)** Hierarchical agglomerative cluster showing the differentiated *shCTRL* from *shTP53* proteomes. **B)** Venn diagram illustrating the total number of significant proteins specifically or commonly regulated by p53 in either untreated or mafosfamide-treated conditions. **C-D)** Volcano plots of significantly regulated proteins after mafosfamide treatment. Each dot represents a protein. Coloured dots indicate significantly up (red) or down (green) regulated proteins.

**Supp.Figure S5. *shCTRL* and *shTP53* hMB cells-derived EVs characterization. A)** Bar plot showing the percentage of DiD-labeled hMB EVs macrophage uptaken by flow cytometry. **B)** Number of 7AAD^+^ hMB cells per µl measured by flow cytometry after treatment with different concentrations of *shTP53-*EVs during 16 or paraphormaldehyde 4% (10 min) as positive control. **C)** Alemtuzumab-mediated ADCP of control hMB cells on the presence of different concentration of *shTP53-*EVs. **B-C)** A concentration of 100 stands for the normal EV concentration used in the ADCP assays described in materials and methods section, and a concentration of 0, indicates that only vehicle (PBS), was added. **D)** Bead-based phagocytosis assay where J774A.1 macrophages were exposed to *shCTRL*-or *shTP53*-EVs 16 h prior to the adding of DyLight680 labeled latex beads.

**Supp. Figure S6. PDL1 expression and function on lymphoma B-cells. A)** Net plot showing PDL1, CD47 and CD200 protein expression change on *shCTRL* and *shTP53* hMB cells after mafosfamdie treatment by flow cytometry (n=3). **B)** Level of ADCP of *Myd88p*.*L252P* cells co-cultured with peritoneal macrophages. All the cells were sublethally pretreated with mafosphamide (1.64 µM) and co-cultures were simultaneously treated with anti-CD20 antibody (50µg/ml) together with either anti-PDL1 (20µg/ml) or anti-PD1 (50µg/ml) for 16hrs (n=3). (*p < 0.05).

