## Supplemental FIgures for "Loss of TP53 mediates suppression of Macrophage Effector Function via Extracellular Vesicles and PDL1 towards Resistance against Chemoimmunotherapy in B-cell malignancies"

### Supplementary Figure S1

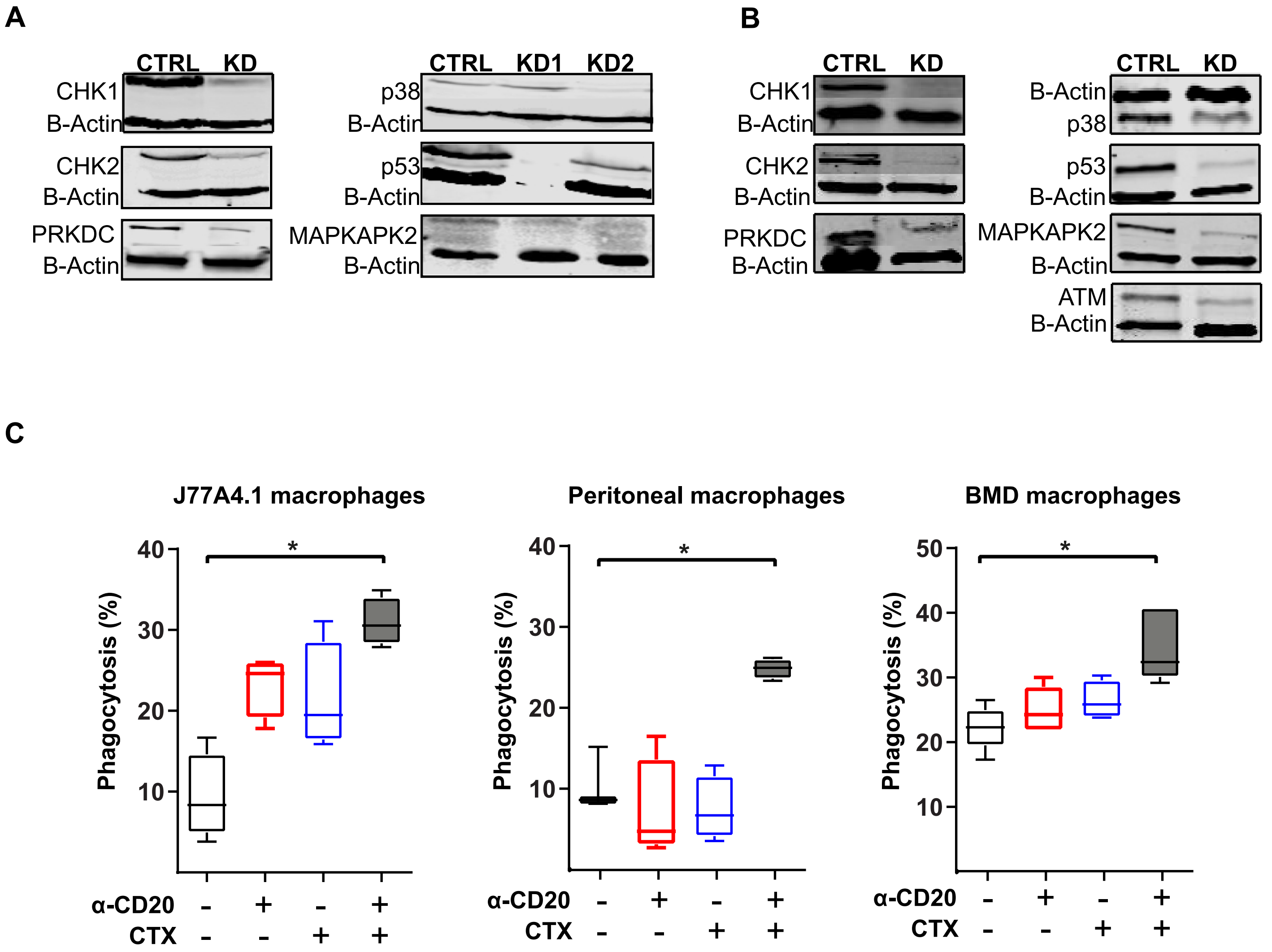

Supplementary Figure S2

A

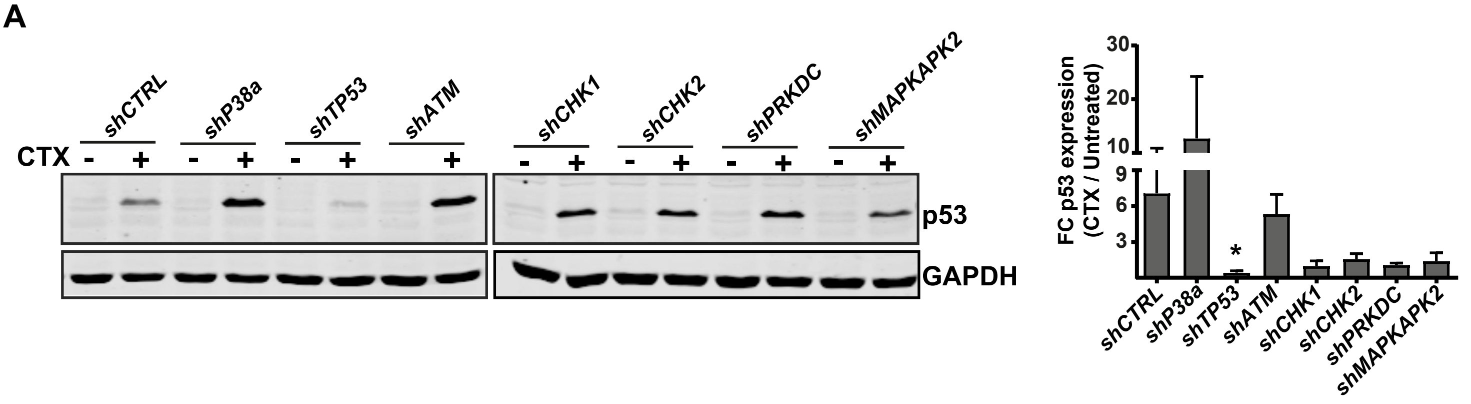


B

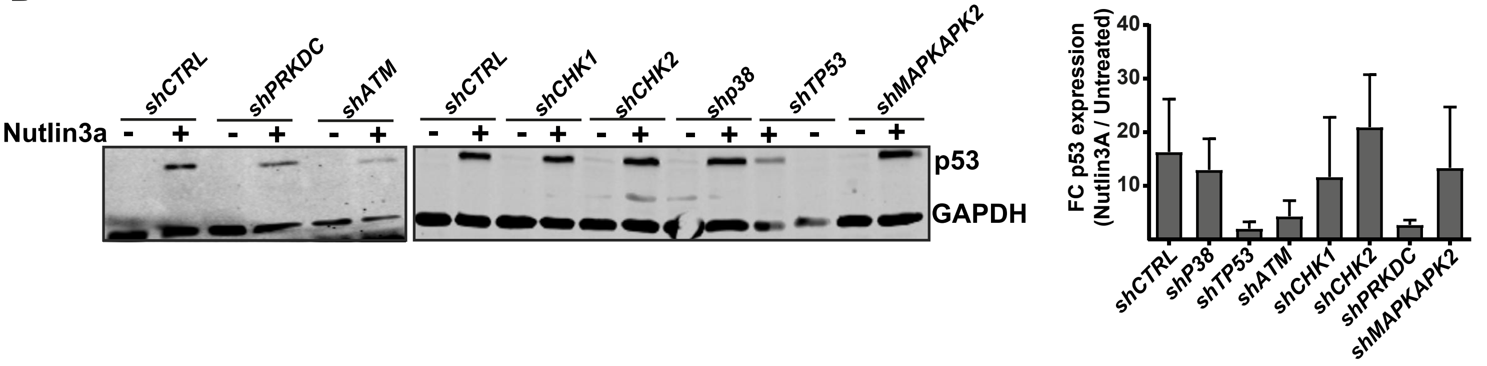

### Supplementary Figure S3

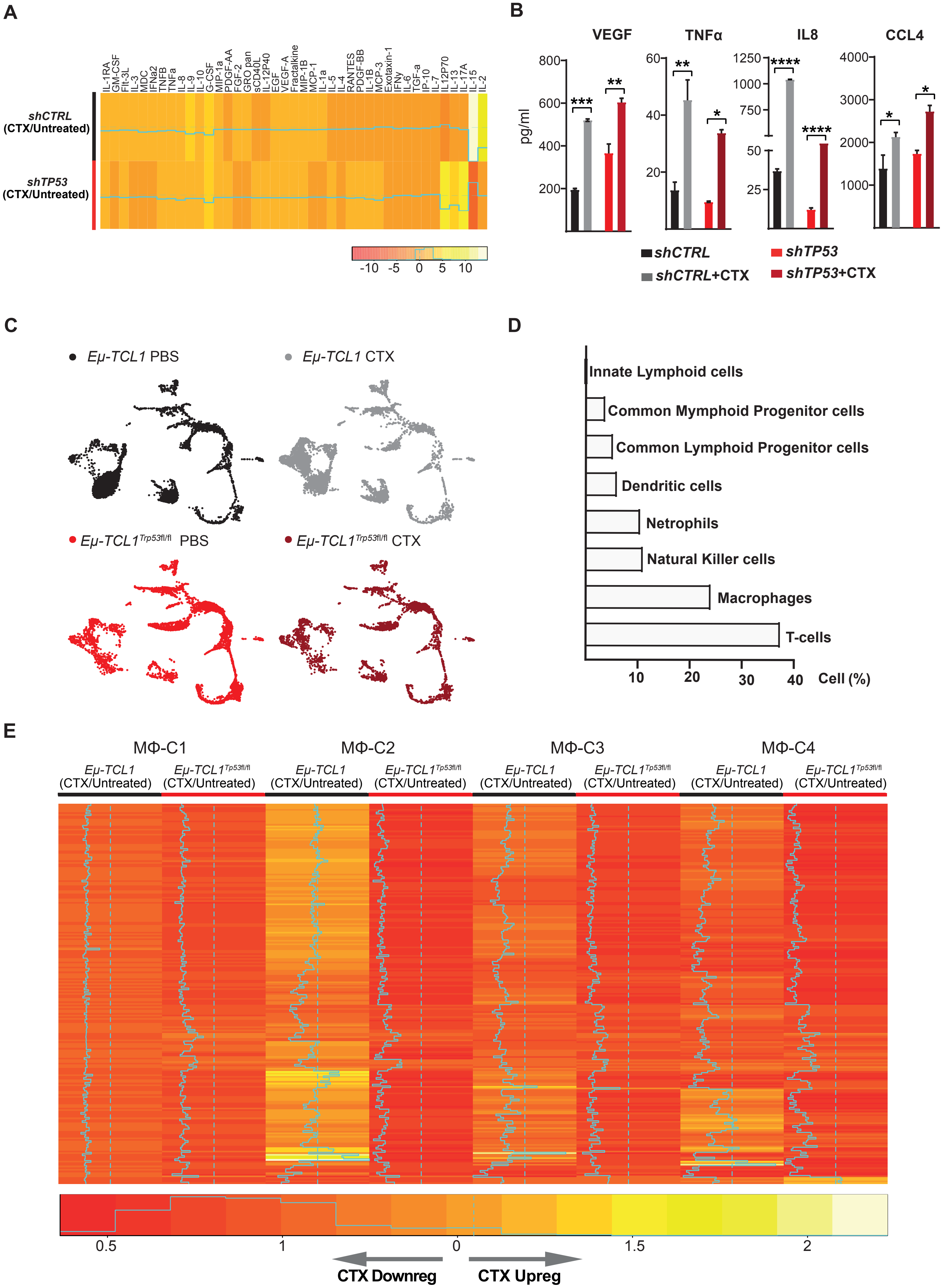

### Supplementary Figure S4

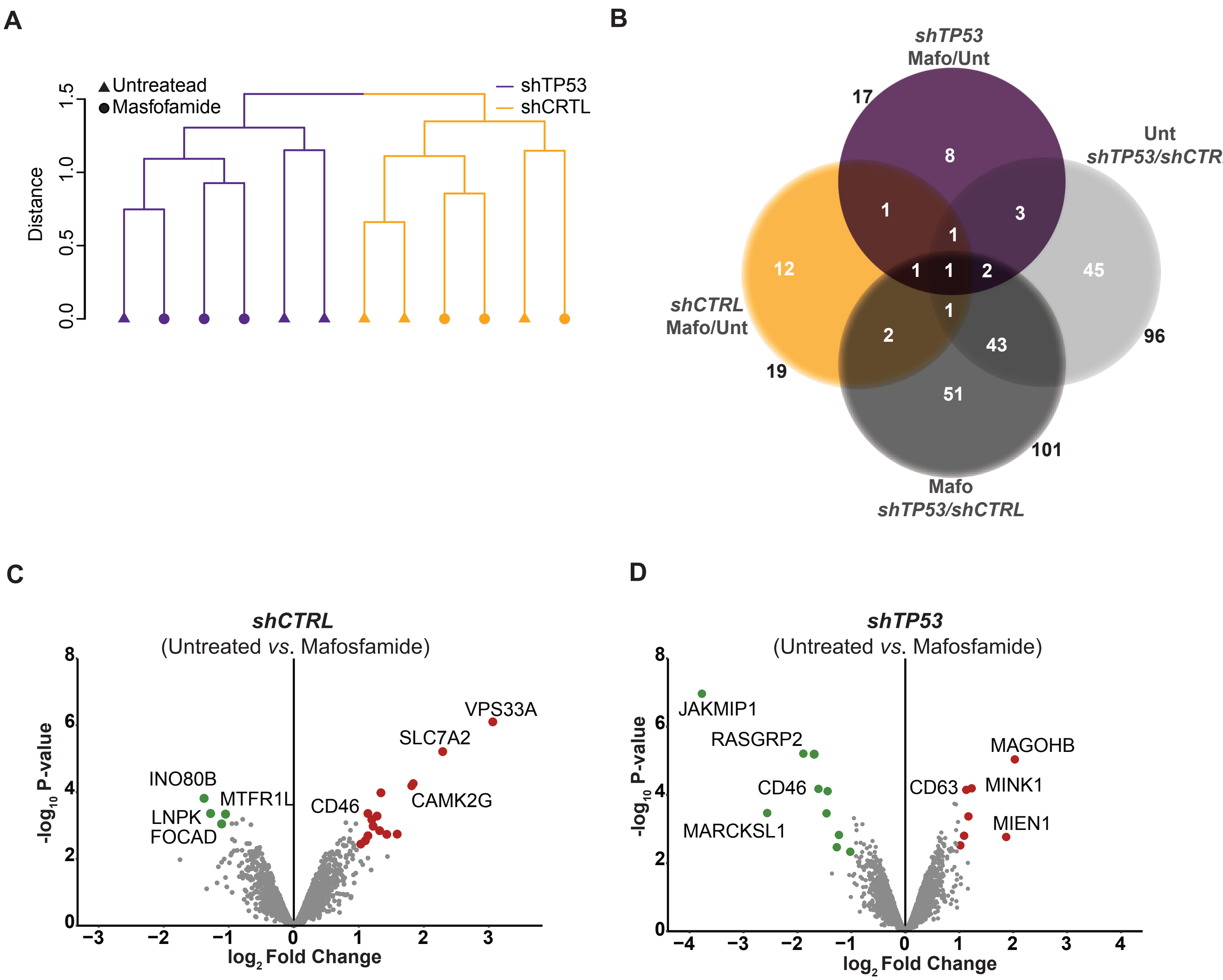

### Supplementary Figure S5

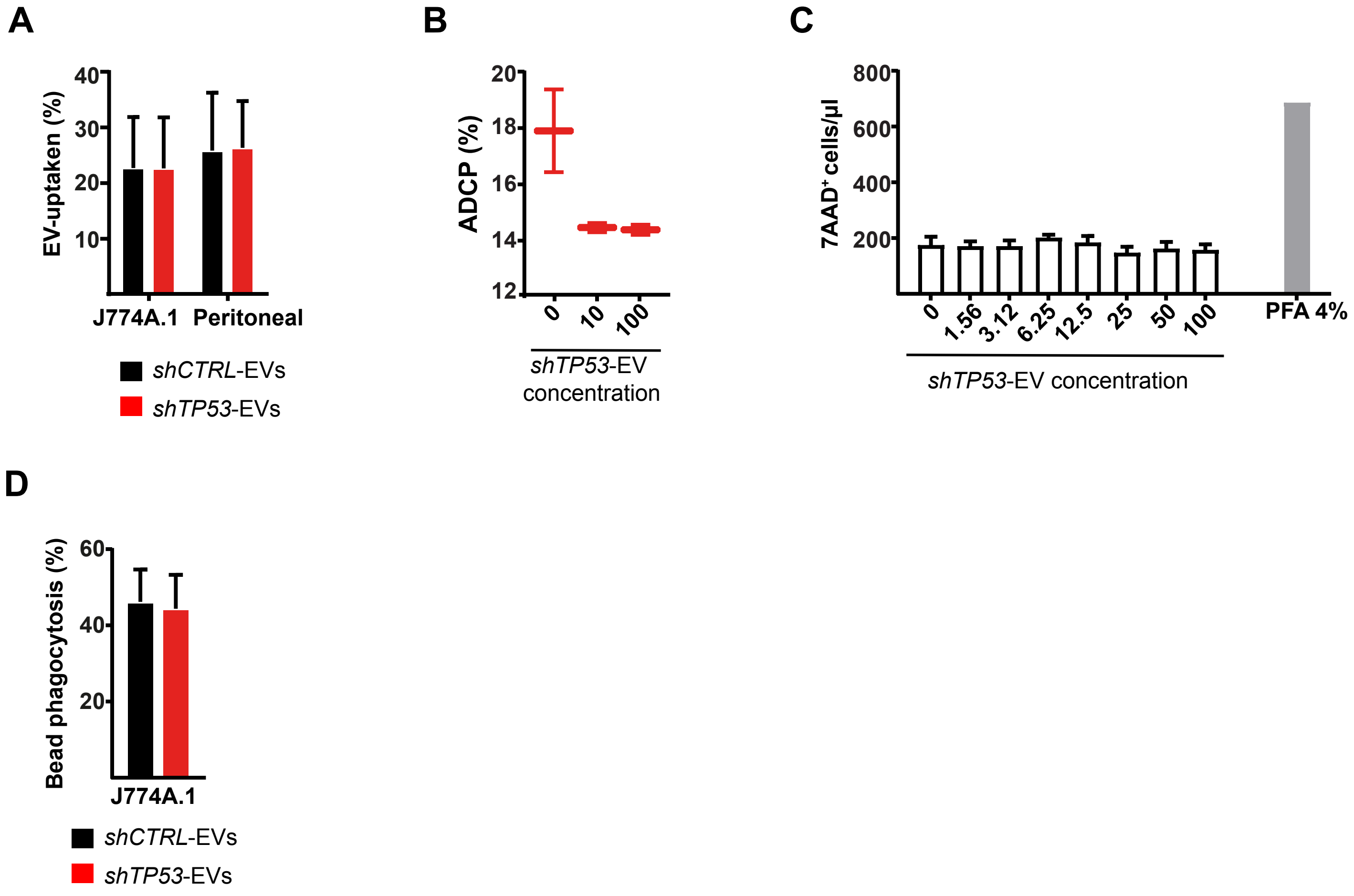

### Supplementary Figure S6

A

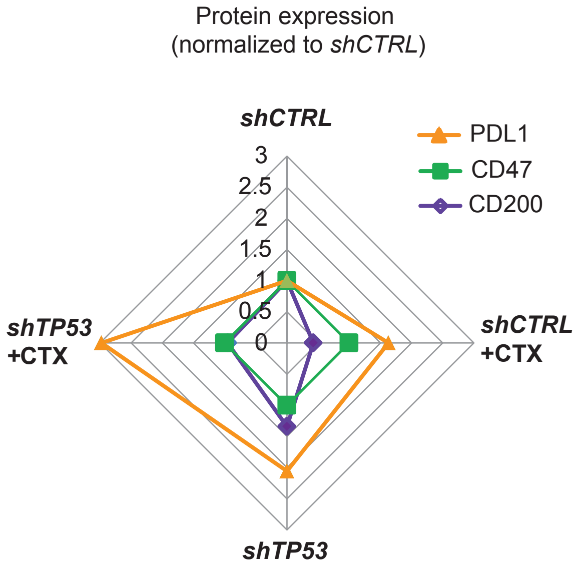


B

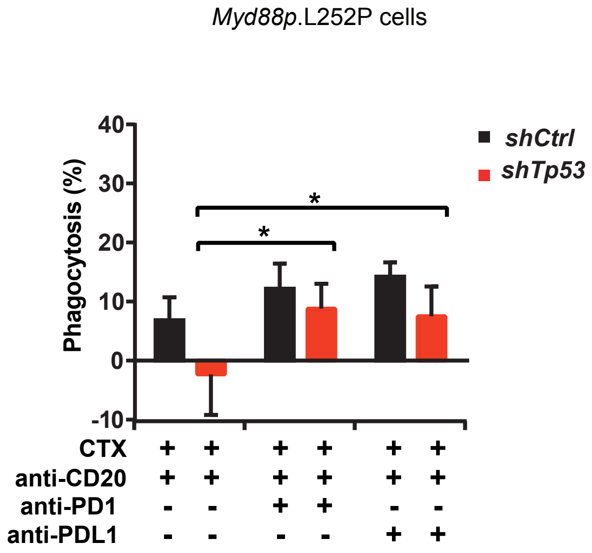


-
